# Cellular and environmental dynamics influence species-specific extents of organelle gene retention

**DOI:** 10.1101/2022.10.17.512581

**Authors:** Belén García Pascual, Jan M. Nordbotten, Iain G. Johnston

**Affiliations:** Department of Mathematics, University of Bergen, Bergen, Norway; Computational Biology Unit, University of Bergen, Bergen, Norway

## Abstract

Mitochondria and plastids rely on many nuclear-encoded genes, but retain small subsets of the genes they need to function in their own organelle DNA (oDNA). Different species retain different numbers of oDNA genes, and the reasons for these differences are not completely understood. Here we use a mathematical model to explore the hypothesis that the energetic demands imposed by an organism’s changing environment influence how many oDNA genes it retains. The model couples the physical biology of cell processes of gene expression and transport to a supply-and-demand model for the environmental dynamics to which an organism is exposed. The tradeoff between fulfilling metabolic and bioenergetic environmental demands, and retaining genetic integrity, is quantified for a generic gene encoded either in oDNA or in nuclear DNA. Species in environments with high-amplitude, intermediate-frequency oscillations are predicted to retain the most organelle genes, whereas those in less dynamic or noisy environments the fewest. We discuss support for, and insight from, these predictions with oDNA data across eukaryotic taxa, including high oDNA gene counts in sessile organisms exposed to day-night and intertidal oscillations (including plants and algae) and low counts in parasites and fungi.

## 1 Introduction

Most eukaryotic cells contain bioenergetic organelles: mitochondria and, in the case of photosynthetic organisms, plastids (including chloroplasts, being a type of plastid). These organelles were originally independent bacteria with their own genomes, and through processes of endosymbioses, they transferred most of their genes to the nuclear genome of the host cell or lost them completely (Martin et al., 2015; Sagan, 1967; Sloan et al., 2018; Adams and Palmer, 2003; Roger et al., 2017; Zaremba-Niedzwiedzka et al., 2017). However, they have retained a small subset of their genes in their own organelle DNA (oDNA), with oDNA in different species containing strikingly different gene counts (Adams and Palmer, 2003; Giannakis et al., 2022; Daley and Whelan, 2005; Johnston and Williams, 2016).

Transfer to the nucleus provides some genetic advantages to organelle genes that are absent in the organelle genome (Sloan et al., 2018; Adams and Palmer, 2003). Nuclear encoding helps avoid sources of mutation in the organelle compartments, which include chemical damage from free radicals (Allen and Raven, 1996) and replicative errors (Kennedy et al., 2013) (which may be more dominant (Itsara et al., 2014)). Nuclear encoding also helps the avoidance of Muller’s ratchet (the irreversible process of accumulation of deleterious mutations) present in the organelles (Daley and Whelan, 2005; Blanchard and Lynch, 2000) and allows sexual recombination and DNA repair in the nucleus (Allen and Raven, 1996). Given these advantages, the question of why bioenergetic organelles retain the genes that they do has been debated for years. At the most fundamental level, there are two pertinent subquestions. The first, gene-centric, question is what makes a given gene more or less likely to be retained in oDNA. The second, species-centric, question is what makes a given species more or less likely to retain a higher or lower number of oDNA genes.

The first question, why a given gene is more or less likely to be retained in oDNA, has had several potential answers proposed over time (summarised in Giannakis et al. (2022)). It has recently been shown that the combination of these can help explain gene-specific patterns of oDNA retention across organelles and eukaryotes (Giannakis et al., 2022). Of particular note here are the hydrophobicity hypothesis (von Heijne, 1986) and the co-localisation for redox regulation (CoRR) hypothesis (Allen and Raven, 1996; Allen, 2015). The hydrophobicity hypothesis asserts that hydrophobic gene products are harder to import to the organelle from the outside (either due to translocation into the organelle (Popot and de Vitry, 1990; Adams and Palmer, 2003) or mistargeting (von Heijne, 1986; Björkholm et al., 2015), proposing that organelle genes encoding hydrophobic gene products are more likely to be retained in the organelles. The CoRR hypothesis proposes that genes are retained in oDNA to allow local, tight control of the energetic machinery (Allen, 2015), so that organelles can better adapt to imposed energetic demands. This idea is supported by the importance of retained oDNA genes in controlling redox processes (Martin et al., 2015; Allen, 2015; Giannakis et al., 2022).

The second question, why some species retain more organelle genes than others, remains more open. There exists substantial diversity in oDNA gene counts across eukaryotes (Roger et al., 2017; Giannakis et al., 2022). Some jakobid protists retain over 60 protein-coding mitochondrial DNA (mtDNA) genes, plants retain fewer and metazoa fewer still with a common 13-protein gene profile shared by a large majority of taxa (including humans). The highest plastid gene counts found so far appear in the group of red alga Rhodophyta with over 200 protein-coding plastid DNA (ptDNA) genes and up to 35 protein-encoding mtDNA genes. In contrast, parasitic species (notably including alveolates) contain very few protein-coding mtDNA genes and some have even lost mtDNA entirely (Hjort et al., 2010). Parasitic organisms can hijack metabolic and energetic budgets of their hosts, so presumably, organelle genes are lost since their bioenergetic organelles have fewer required functions, but how can the diversity of oDNA gene counts in other taxa be explained?

To address this species-specific question, we focus on how the energetic demand imposed by the environment changes over time. Following the CoRR idea that retaining genes in oDNA improves organelles’ local responses to changing conditions, we hypothesise that organisms facing large and/or rapid environmental changes in bioenergetic demand require more local control over organelle machinery to respond to these changes (Johnston, 2019a). The hypothesis suggests that organisms in stable and low-demand environments (including parasites) require less organelle control, as the production of energy is not so challenging. Organisms subject to more dynamic environmental demands (for example, the diurnal oscillations of light, or semidiurnal oscillations of tide) require tighter control over challenging energetic and metabolic demands, so these organisms are predicted to retain a higher count of oDNA genes.

Hypotheses about evolutionary processes can be challenging to test with experiments. Quantitative modelling can shed light on evolutionary pressures and dynamics, as demonstrated by powerful theoretical studies exploring the co-evolution of oDNA and the host cell (Radzvilavicius et al., 2016, 2017, 2021; Hadjivasiliou et al., 2012, 2013), and models of the specific features involved in oDNA gene retention (Kelly, 2021; Giannakis et al., 2022), although few quantitative models to our knowledge have explored CoRR in the same depth. Here, to test our hypothesis, we propose and analyse a mathematical model for a given organelle gene in a given organism. This model captures the interplay of cellular processes (like gene expression, import of gene product to the organelle and degradation), environmental dynamics (like the amplitude and frequency of an environmental wave, also understood as the energetic demand placed on the organism) and the proportion of wild-type organelle DNA which allows the organelle to functionally synthesise the gene products it needs. We use the model to probe how environmental dynamics influence the organism’s ability to meet energetic demands for each of the two cases of encoding compartments: the nucleus and the organelle, providing a general quantitative framework to explore links between CoRR, environmental dynamics and gene retention.

## 2 Methods

### 2.1 Model description

The mathematical model explores the ability of an organism to meet environmental demands, considering the interplay of cell biological processes and particular model environments. It is summarised in Figure 1 and built up in stages as follows.

**Figure 1:**
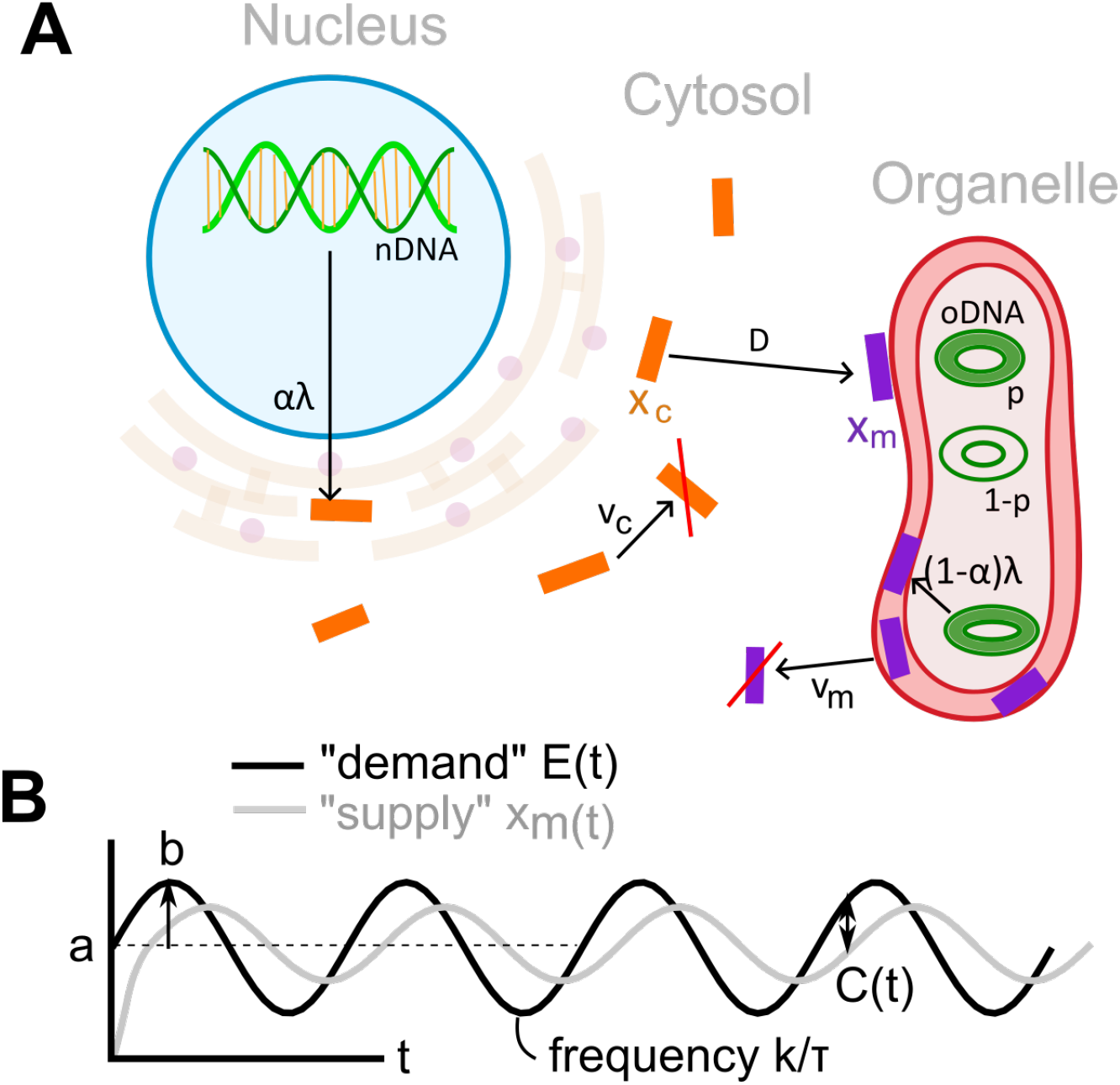
Model outline. **(A)** The cell biological processes modelled in gene expression and translocation. A gene encoded in the organelle (*α* = 0) is expressed with rate *λ* from wild-type oDNA, whose proportion is *p*. An organelle gene encoded in the nucleus (*α* = 1) is expressed from nDNA with rate *λ*, and its gene product is imported to the organelle with a transport rate *D*. The labels *x*_*c*_ and *x*_*m*_ represent the amount of gene product in the cytosol and organelle, respectively, and *ν*_*c*_ and *ν*_*m*_ are degradation rates in those compartments. **(B)** Environmental supply and demand. The environment places demand on the organelle machinery. For oscillating signals this demand has mean *a*, relative amplitude *b* and frequency *k/τ*. Organelle gene products *x*_*m*_ will respond to this demand: following some transient behaviour an oscillation will also occur. An instantaneous cost *C*(*t*) is incurred, being the absolute difference between environmental demand *E*(*t*) and functional machinery supply *x*_*m*_(*t*).

#### 2.1.1 Cell biological processes: gene expression, transport and degradation

The set up of the model is for a generic gene encoding organelle machinery. The gene may be encoded in the organelle or in the nucleus of the cell, in which case it must be transported to the organelle through the cytosol. Working at a coarse-grained level, we will use this general picture to describe both mitochondria and plastids. The model has two dynamic variables: *x*_*m*_(*t*) is the available amount of functional gene product in the organelle at time *t* (for example, the various protein subunits of electron transport chain complexes) and *x*_*c*_(*t*) is the available amount of gene product in the cytosol at time *t*.

The parameters of the model for the cellular processes are the following non-negative constant rates: *λ* is the synthesis rate of gene product (which will be scaled by the cell’s response to the environment, see below), *D* is the import or transport rate of gene product from the cytosol to the organelle, *ν*_*m*_, *ν*_*c*_ are the degradation rates of gene product in the organelle and in the cytosol, respectively, and *p* is the proportion of wild-type oDNA, also called the genetic integrity, which allows the synthesis of functional gene product in the organelle. The cell processes whose rates are the ones described above are illustrated in Figure 1A.

The parameters *D* and *ν*_*c*_ describe the ability of a nuclear-encoded gene product to translocate to its required position in the organelle. These can be used to model different mechanisms proposed for the hydrophobicity hypothesis. One mechanism is that hydrophobic gene products cannot readily be unpacked to import into the organelle (Popot and de Vitry, 1990; Adams and Palmer, 2003), corresponding to a high value of the degradation rate in the cytosol *ν*_*c*_, as the gene product is merely lost and hence can be considered as degraded. Another mechanism is that the gene product is mistargeted, usually to the endoplasmic reticulum (von Heijne, 1986; Björkholm et al., 2015), so that the import to the organelle takes a much longer time: corresponding to a low value of the transport rate *D*.

#### 2.1.2 Coupling of the environmental supply-and-demand model and the cell biology processes

We denote by *E*(*t*) the energetic demand placed on the cell by the environment. The goal of the cell is to match *x*_*m*_(*t*), the ‘supply’ of gene product present in the organelle, to environmental demand *E*(*t*). We picture the cell as sensing, and able to respond to, environmental demands by expressing genes that support organelle function. Hence we define a feedback signalling function *f* (*E*(*t*), *x*_*m*_(*t*)) that controls the production of gene product as

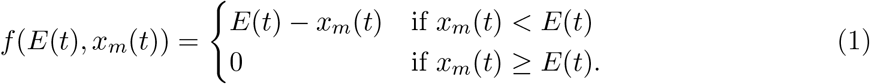

Since the dynamics depend on the compartment where the organelle gene is encoded (the organelle itself or the nucleus), we use a coefficient *α* as a model index representing the encoding compartment, where *α* = 0 if the gene is in the organelle or *α* = 1 if the gene is in the nucleus.

We then have our model of first-order, linear ordinary differential equations describing the instantaneous rates of change of the variables *x*_*m*_(*t*) and *x*_*c*_(*t*) as follows:

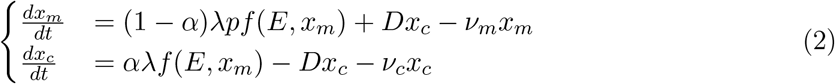

for either *α* = 0 or *α* = 1. We take initial conditions (*x*_*m*_(0), *x*_*c*_(0)) = (0, 0), but we focus on long-term behaviour after the corresponding transient periods have disappeared, described below.

The mathematical properties of Eq (2) are studied in the appendixes (Supplementary Information), including well-posedness (A.1) together with existence and stability of equilibrium points (A.2 and A.3).

#### 2.1.3 Bioenergetic cost

The model is of supply-and-demand nature by Eq (1), so as in previous quantitative work (Hoitzing et al., 2019), we can define a cost function that measures how well the organelle supplies the organism with the energy that its environment demands. A high cost corresponds to supply far away from the demand, either surplus or deficit. This cost function is for a specific encoding compartment, so that the compartment incurring the lowest cost is interpreted as the most favourable in which to encode the gene. We define the cost function as the absolute difference between environmental demand *E*(*t*) and supply of functional gene product in the organelle *x*_*m*_(*t*) integrated over the time window [*t*_*i*_, *t*_*f*_]:

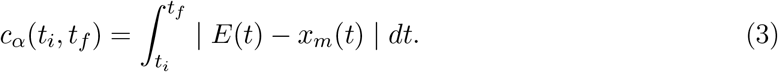

This supply-and-demand model is illustrated in Figure 1B.

To ensure the robustness of our results we consider different choices of cost functions in the Supplemental Information (Figures S3, S12 and S13) and show that the model’s behaviour is similar across several choices.

#### 2.1.4 Specific types of environmental demands

We explore the extent of oDNA genes that an organisms is most likely to retain by looking at the influences that different types of environmental demands *E*(*t*) have on the system. We consider static environments, periodically changing environments and randomly changing environments. Static environments are simply modelled with *E*(*t*) = *a*. For the case of a periodically changing environment, let *a >* 0 be the time-averaged value of environmental demand, *ab* the amplitude of the oscillation (with *b* seen as the relative amplitude), *τ* the characteristic time-scale of the system (for example a day in a physical context) and *k* the frequency of the oscillation relative to the time-scale *τ* (Figure 1 (B)). We then model the energetic demand placed on the organism by a periodically changing environment as a wave *E*(*t*) = *E*_*p*_(*t*) where *E*_*p*_(*t*) is defined as:

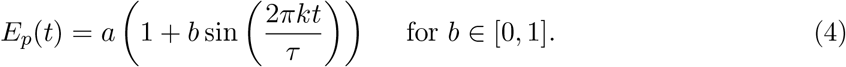

For a randomly changing environment, we consider *E*(*t*) as a stochastic process which is either uncorrelated white noise (as used in descriptions of terrestrial environments (Vasseur and Yodzis, 2004)), or correlated red noise (as used in descriptions of marine and coastal environments (Vasseur and Yodzis, 2004)), also called a random walk.

For numerical implementation, for a time interval [0, Γ] of the model, we define a partition 0 = *s*_0_ *< s*_1_ *<* … *< s*_*k*−1_ *< s*_*k*_ = Γ so that *D* = {*d*_*i*_}_*i*∈*I*_, where *d*_*i*_ = [*s*_*i*_, *s*_*i*+1_], is the collection of time subintervals. We then consider *D* to be the time-discrete, finite and bounded domain of the time-continuous series of white and red noise. The range of both types of noise is [0, 2*a*], so that the average value is *a* as for the periodically changing environment Eq (4).

For each *i*, we draw a random number *u*_*i*_ that has a uniform distribution in [0, 2*a*], i.e. *u*_*i*_ ∼ *U* (0, 2*a*). White noise *E*_*w*_(*t*) is then defined as:

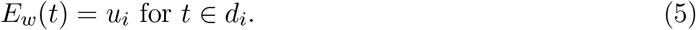

To define red noise *E*_*r*_(*t*), we write the accumulation of *u*_*i*_ with a step-size *d*_*i*_ = *s*_*i*+1_ − *s*_*i*_, and we keep it bounded in [0, 2*a*] as follows:

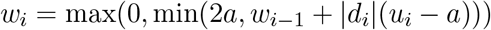

then we define *E*_*r*_(*t*) as:

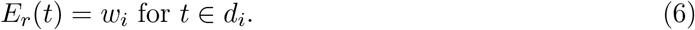

### 2.2 Biological parameterisations

The specific values of the rates involved in our model vary substantially across species, and across genes in the same species. However, we can connect with biological quantities at a coarse-grained level to interpret our model in an informative way (Johnston et al., 2014).

We take characteristic rates measured for the case of yeast *Saccharomyces cerevisiae* as reference for our model. The synthesis rate of gene product is a combination of transcription and translation, the rates of which vary substantially across genes; we will explore orders-of-magnitude ranges in these parameters throughout this study. As first estimates, a transcription rate of 0.12 min^−1^ message RNA molecules (Pelechano et al., 2010) and a translation rate of 0.43 min^−1^ protein molecules (Siwiak and Zielenkiewicz, l 29) give an overall rate of around 0.1 min^−1^. Therefore, we choose *T* = 10 min to be the time unit in our model, so that the synthesis rate is set as *λ* = 1*T* ^−1^ for simulations by default (the effect of different synthesis rates are shown in Figures S10 and S11 in the Supplementary Information). The average scale of protein half-lives in the cytosol of 43 min (Belle et al., 2006) give *ν*_*c*_ ≈ 0.1*T* ^−1^. Protein half-lives in the mitochondrion vary dramatically across hours and days for different genes, suggesting a possible range of *ν*_*m*_ ≈ 0.01 −0.1*T* ^−1^ (Christiano et al., 2014). Import rates to the mitochondrion also vary substantially across genes, suggesting *D* ≈ 0.1 −10*T* ^−1^ (Schäfer et al., 2022). The model parameters that we map therefore contain a range of values that correspond to plausible dynamics for real genes. Different eukaryotic species will have different values for these parameters, but the ranges we consider are compatible with observations across kingdoms (for example, Li et al. (2017)).

We set a time window [0, 2 × 144] of two days in *T* time units for the numerical simulations. To remove the transient periods determined by the initial conditions (*x*_*m*_(0), *x*_*c*_(0)) = (0, 0) and focus on the equilibrium phases of the system, we use the first day [0, 144] as an equilibration step. Then we take the solutions of the model and compute the respective cost functions Eq (3) only for the last full day, which corresponds to the time window [144, 2 × 144].

In the periodic environment *E*_*p*_(*t*) Eq (4), we set the characteristic time-scale *τ* to be a full 24 hours period in *T* units, i.e. *τ* = 144*T* (*τ* = 144*T* = 144 × 10 min = 24 hours). So *k* = 1 corresponds to diurnal oscillation (as with day-night cycles), and *k* = 2 to semidiurnal oscillation (as with tides).

The wild-type proportion of oDNA *p* is a coarse-grained measure of oDNA integrity in our model. Lower *p* corresponds to more oDNA damage, compromising the expression of functional organelle machinery; *p* = 1 corresponds to perfect oDNA and hence the maximum possible expression capacity. The propensity for oDNA damage in an organism, both within a lifetime and across generations, will act to reduce *p*. Rather than attempting to set this value to match a given biological instance, we will explore the behaviour of the system as it varies. Furthermore, *p* can be pictured as a snapshot value from the ongoing process of oDNA damage accumulation, segregation, and mitigation (Johnston, 2019b; Guo et al., 2013).

### 2.3 Numerical implementation

The numerical simulations and visualisations of results were all performed in Python. The model of ordinary differential equations presented in Eq (2) was solved using the numerical integrator scipy.integrate.odeint that uses the method LSODA from the FORTRAN library odepack (Hindmarsh, 1983; Radhakrishnan and Hindmarsh, 1993). The heatmaps were made using seaborn (Waskom, 2021) and the time-lines and phase portraits were made using matplotlib.pyplot (Hunter, 2007). The integral of the cost function Eq (3) was computed using the composite Simpson’s rule using scipy.integrate.simps. Mathematical expressions were implemented using NumPy (Harris et al., 2020) and mpmath (Johansson et al., 2013). The code can be found in 0https://github.com/StochasticBiology/Environmental-oDNA-retention.git.

### 2.4 Bioinformatics

The curation of oDNA gene counts follows and builds upon the pipeline for eukaryotic oDNA analysis in (Giannakis et al., 2022). All available complete mtDNA and ptDNA sequences were downloaded from RefSeq (O’Leary et al., 2015). Gene annotations were systematised with BioPython (Cock et al., 2009) according to a manually-curated list of label substitutions, taken from and validated in (Giannakis et al., 2022). The subset of protein-coding genes in these lists (omitting RNA genes, gene fragments, open reading frames not known to encode a protein, and various other anomalous entries) present in each species’ oDNA was then recorded. The species in the dataset were embedded in a Common Taxonomy Tree (Federhen, 2012) and the members of each basal eukaryotic clade compiled into a corresponding set using R (R Core Team, 2020), with libraries ggplot2 (Wickham, 2016), gridExtra (Auguie, 2017), and phytools (Revell, 2012).

## 3 Results

Our model, described in Methods, pictures the environment as imposing metabolic and bioenergetic demands on the cell, which may be static or vary periodically or randomly over time. This may correspond, for example, to diurnal variation in light levels, temperature, animal activity, and so on; to semidiurnal tidal variation in oxygenation and salinity; or to more rapid variation due to bursts of activity, fluctuating shade, or other conditions. The cell expresses organelle genes to produce gene products in an attempt to supply the required machinery to meet this demand. Genes may be encoded in either the nucleus, in which case their gene products must be imported to the organelle, or in the organelle, in which case the genes are potentially subject to mutational damage. We integrate the absolute difference between supply and demand over a simulated time window to calculate the cost for each compartment Eq (3) for a given parameterisation.

There are several parameters in the model corresponding to cell biological and environmental quantities (Milo and Phillips, 2015). Some key parameters are the synthesis of gene product *λ*, the transport *D* and the degradation of gene product in the organelle *ν*_*m*_ and in the cytosol *ν*_*c*_. We also consider a measure of oDNA integrity, interpreted as the proportion *p* of wild-type, functional oDNA. Rather than attempt an exhaustive characterisation of the full parameter space in the main text, we focus on several cases which illustrate the more general trends, and support these examples with other results in the Supplementary Information (SI). We explore biologically plausible ranges of parameters, with simulation time unit *T* = 10 min setting the scale for other parameters (see Methods), and simulating for a time-scale of a full day after allowing transient behaviours to disappear.

### 3.1 Static environments

To focus on the influence that the cell biological parameters (see Methods), have in the model, we first look at the case of a static environment *E*(*t*) = *a* for all *t*.

In Appendix A.2 (SI) we algebraically find the solutions of the system when it is in equilibrium, and from there we can directly compute the instantaneous cost *C*_*α*_ for encoding compartment *α* as the absolute difference between the energetic demand *E*(*t*) = *a* and the supply of gene product in the organelle *x*_*m*_(*t*). We look at the ratio between the instantaneous cost for the organelle-encoding strategy (*α* = 0) and the nuclear-encoding strategy (*α* = 1):

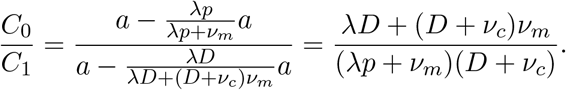

Here, if *C*_0_*/C*_1_ *>* 1, then encoding the gene in the nucleus is the most favourable strategy, and if *C*_0_*/C*_1_ *<* 1, then encoding in the organelle is the most favourable strategy. The first condition *C*_0_*/C*_1_ *>* 1 holds if *D > p*(*D* + *ν*_*c*_), and this is the case when *D* is high (the import to the organelle is fast), *ν*_*c*_ is low (gene product is not lost in the cytosol so the import to the organelle is more efficient) and *p* is low (the organelle has a significant load of mutant oDNA that prevents the synthesis of gene product). The second condition *C*_0_*/C*_1_ *<* 1 holds if *D < p*(*D*+*ν*_*c*_), and this is the case when *D* is low (the import to the organelle is slow), *ν*_*c*_ is high (a lot of gene product is degraded and therefore not so much is being imported to the organelle) and *p* is high (there is good proportion of wild-type oDNA to synthesise gene product). If we have the special case of no degradation in the organelle *ν*_*m*_ = 0, then *C*_0_*/C*_1_ = 1, meaning that both strategies are equally favourable. These trends are clearly observable in the numerical results in Figure 2A, illustrating a moderate level of oDNA damage (*p* = 0.75). Without oDNA damage (*p* = 1), there is never a reason to favour the nuclear compartment, and nuclear-encoded costs at best match (but never drop below) organelle-encoded costs. Figure S2 in the SI explores higher values of *p*.

**Figure 2:**
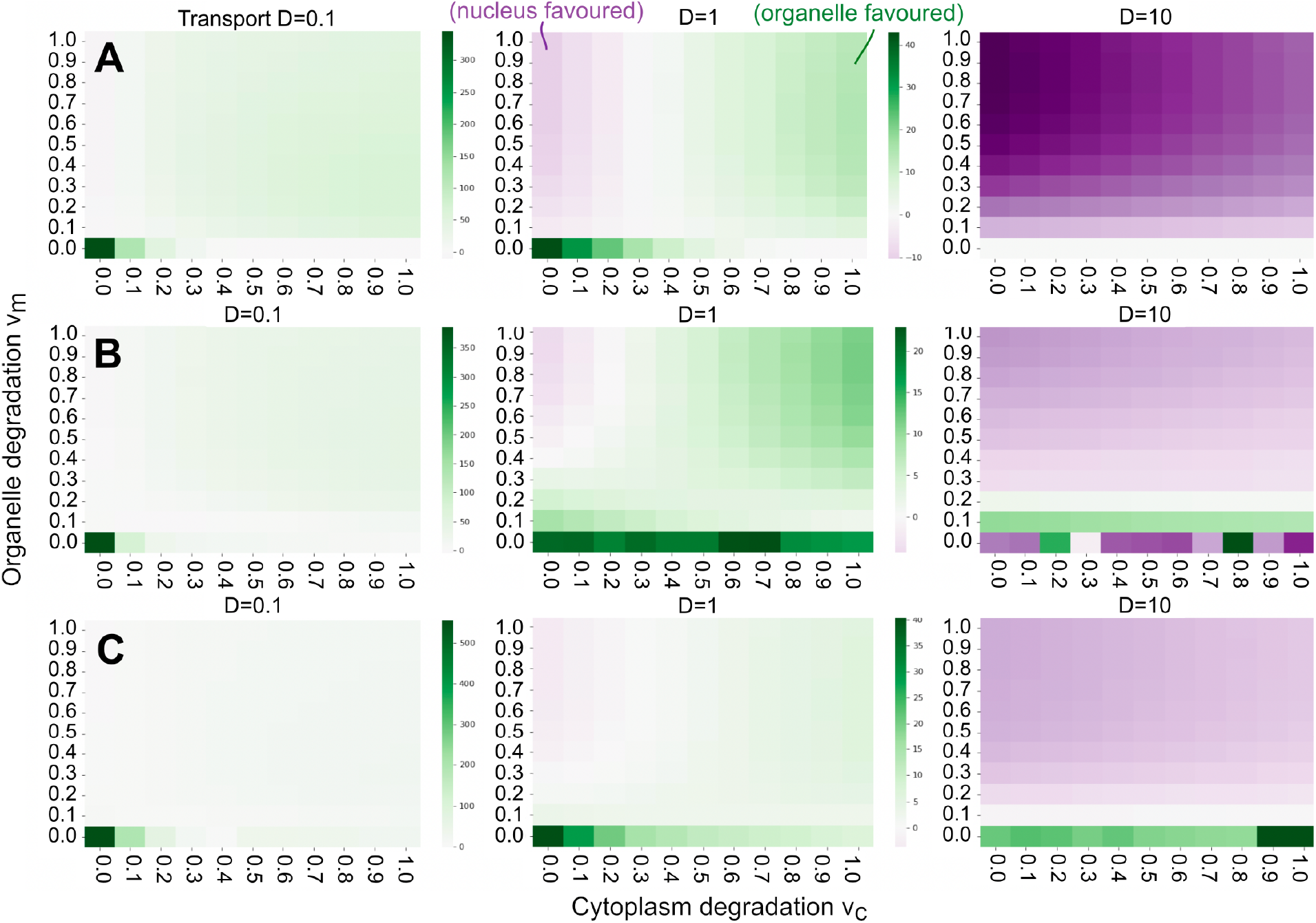
Cost differences with the interplay of degradation and transport in static and noisy environments. Absolute cost difference between nuclear and organelle encoding strategies for a **(A)** static *E*(*t*) = *a*, **(B)** white noise *E*_*w*_(*t*) Eq (5) and **(C)** red noise *E*_*r*_(*t*) Eq (6) environment. The parameter values are synthesis rate *λ* = 1, constant environment *E*(*t*) = *a* = 1 and proportion of wild-type oDNA *p* = 0.75. This proportion *p* influences which compartment is more favourable; results for *p* = 1 and *p* = 0.9 for a static environment are in Figure S2 in the SI.

Here and throughout, the parameters of the model provide a way of connecting the theory to specific genes. Gene products with high turnover correspond to a high *ν*_*m*_; long-lived products have a low *ν*_*m*_. Gene products prone to mistargeting (which could include hydrophobic products under the ‘mistargeting’ picture (Björkholm et al., 2015)) have a low *D*; those challenging to import to the organelle (which could include hydrophobic products under the ‘unfolding’ picture (von Heijne, 1986)) have a high *ν*_*c*_. A value of *p <* 1 corresponds to the expectation that genes encoded in oDNA will accumulate some level of genetic damage over time; lower *p* corresponds to more damage, while higher *p* corresponds to more robust oDNA maintenance (Johnston and Burgstaller, 2019).

Degradation rates play an important role in the model. The degradation rate in the organelle *ν*_*m*_ limits the ability of the cell to meet environmental demand: the organelle is prevented from having as much gene product as it needs, and the variable *x*_*m*_(*t*) is always damped with respect to an environmental function *E*(*t*). For low *ν*_*m*_ (long-lived gene products), no significant amount of gene product is being lost and the total amount in the organelle is more correctly modulated by the signalling function Eq (1). For *ν*_*m*_ = 0 the system can perfectly satisfy environmental demand (Appendix A.4 in the SI).

### 3.2 Randomly changing environments

Here we analyse our model for randomly varying energetic demands (see Methods). The environments behave either like uncorrelated white noise Eq (5) or like correlated red noise Eq (6). We see in Figure 2 that the plots for static, uncorrelated and correlated noisy environments show very similar patterns, so we can infer that the same principles for static environmental demands discussed in Section 3.1 also hold for noisy ones. This is because the system tends to adapt to an average environmental demand (either a constant for uncorrelated noise or a moving average for red noise) and fluctuations around this adapted average challenge the system in similar ways.

In Figure S4 in the SI we see that when there is some degradation in the organelle, *ν*_*m*_ = 0.5, and no degradation in the cytosol, *ν*_*c*_ = 0, for uncorrelated white noise the costs for both encoding compartments are almost identical. For correlated red noise, the cost for the nuclear-encoding compartment is lower. These examples agree with the correspondent points in the heatmaps in Figure 2 for *D* = 0.1 and *D* = 1.

### 3.3 Periodically changing environments

Next, we consider oscillating environmental demands as defined in Eq (4). For different frequencies *k* and time-averaged demands *a*, we look at when the amplitude takes a maximal value (*b* = 1), representing higher-magnitude environment oscillations, and when the amplitude is lower (*b* = 0.5), representing an environment closer to the constant case. Qualitatively, the higher the relative amplitude *b* and the frequency *k*, the more dramatically changing the environment is. We see in Figure 3 that encoding the organelle gene in the organelle is much more favourable the more the environment changes, that is, for high amplitude *b* = 1 and every frequency when the transport is slow (*D* = 0.01, corresponding to hard-to-import gene products), and for high frequencies when the transport is faster (*D* = 0.1 and *D* = 1). This agrees with the hypothesis that organisms tend to retain more genes in their oDNA if their environments dramatically and rapidly change, and suggests that those harder to import (possibly including hydrophobic products (Björkholm et al., 2015; von Heijne, 1986)) will be the most retained.

**Figure 3:**
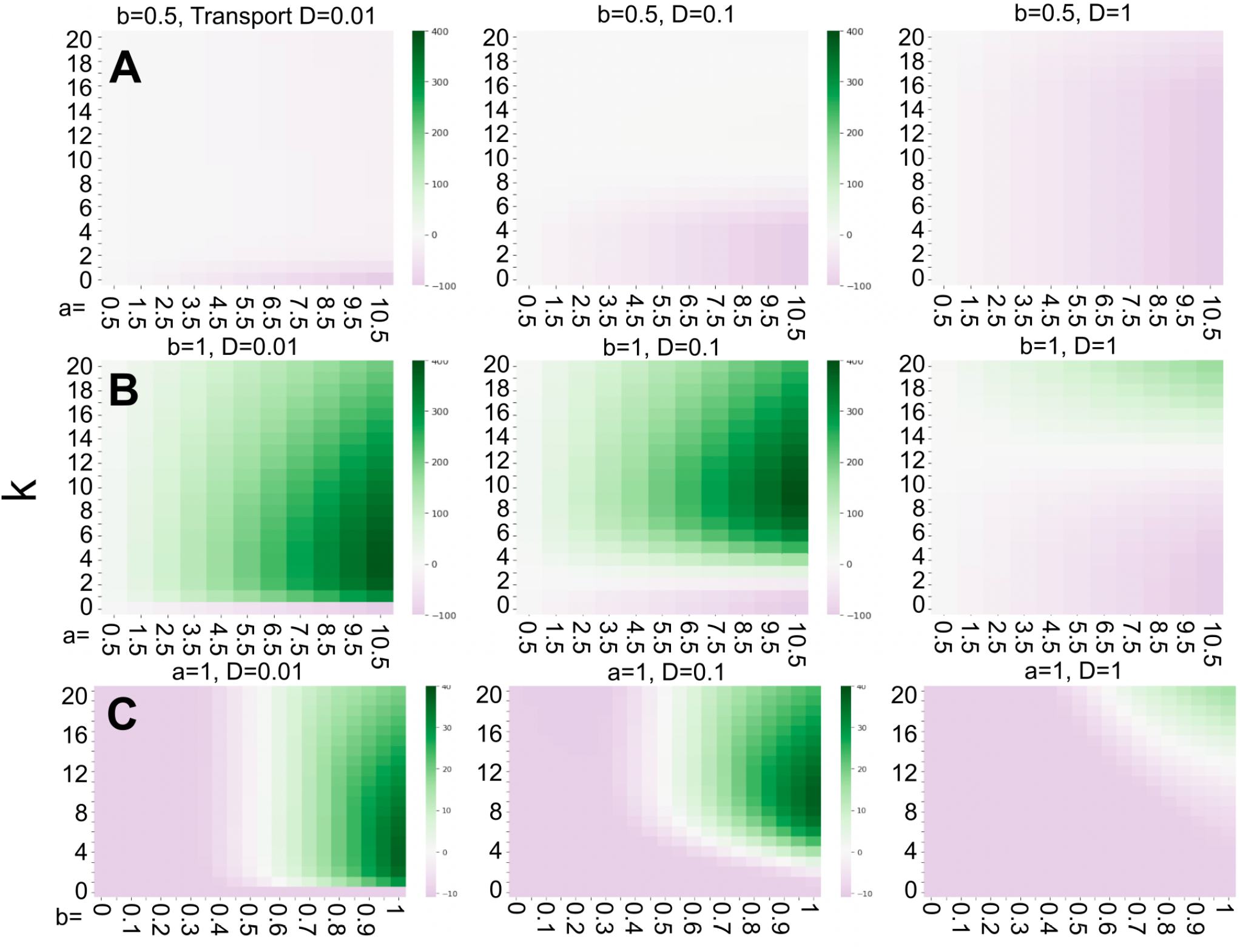
Cost difference with the interplay between environmental oscillation and cell biological processes. Environmental oscillations have frequency *k* per day, amplitude *b* and mean value *a*. Other parameter values are synthesis rate *λ* = 1, degradation in the organelle *ν*_*m*_ = 0.5, no degradation in the cytosol *ν*_*c*_ = 0 and proportion of wild-type oDNA *p* = 0.75. Different parameterisations show largely the same trends and are explored in the Supplementary Information: alternative values of *p* (Figures S5 and S6); nonzero degradation in the cytosol *ν*_*c*_ = 1 (Figures S7 and S9); higher synthesis rate *λ* = 10 (Figures S8 and S11); and low synthesis rate *λ* = 0.1 (Figure S10).

Strikingly, organelle encoding is most strongly favoured at intermediate frequencies *k* of environmental fluctuation, as seen in Figure 3. The specific values of these frequencies depend on the transport rate: for lower transport rates, the peak value for *k* is lower, and for higher transport rates it is higher. Whether this translates to an overall favouring of the organelle compartment also depends on cytoplasmic degradation and oDNA wild-type proportion.

On the other side, the nuclear-encoding case is most strongly favourable for a less dynamic environment. We see in Figure 3 that this compartment begins to dominate for lower amplitude *b* = 0.5, and is completely dominant under these parameterisations for a constant environment (*k* = 0) if the proportion of wild-type oDNA *p* is less than 1. For the case when *p* = 1 (perfect oDNA integrity can be retained), we see in Figures S5 and S6 in the SI that the nuclear-encoding case is never favoured, by the reasons described above. Following the trends observed in the static environment, decreased organelle damage *p* = 1 (Figures S5 and S6 in the SI), or increased cytoplasmic degradation *ν*_*c*_ = 1 (Figures S7 and S9 in the SI), has the straightforward effect of shifting the trends in Figure 3 quite uniformly across parameterisations to favour organelle encoding.

We can again use the parameterisations of the model to consider predicted gene-to-gene behaviour. As an illustration, consider three gene products: one hydrophobic and challenging to import (with *D* = 0.01), one intermediate (with *D* = 0.1) and a less hydrophobic product with less import difficulty (with *D* = 1). Consider also two sets of environmental conditions, *k* = 4 (intermediate frequency) and *k* = 10 (high frequency) with the same *b*. Under these conditions and the other biologically plausible parameters in Figure 3, for intermediate frequency *k* = 4 only the most hydrophobic product (for *D* = 0.01) is favoured in the organelle with the others (for *D* = 0.01 and *D* = 1) favoured in the nucleus. For *k* = 10 only the most hydrophilic (*D* = 1) will remain in the nucleus. Figure S5 in the SI also supports this picture for different proportions of wild-type oDNA *p*.

Lastly, from the algebraic solutions of the model in Appendix A.4 in the SI, we can see explicitly how an oscillatory environment challenges the system, as there is an absence of a perfectly synchronised solution *x*_*m*_(*t*) to *E*_*p*_(*t*) Eq (4). This is because for both organelle-encoding and nuclear-encoding cases the only way the system can reach the time-averaged demand *a* in *E*_*p*_(*t*) is if *ν*_*m*_ = 0, but then it cannot enter to decay phase when the environmental wave *E*_*p*_(*t*) goes down. This scenario places the system out-of-phase with respect to the environmental wave, and therefore leads to a high cost. This is seen in Figure 4 and Figure S1 (vi) in the SI. Moreover, for the organelle-encoding case, the variable *x*_*m*_ cannot match the fluctuating environment simultaneously in phase and in amplitude, as discussed in Appendix A.4. For the nuclear-encoding case, a solution in phase with environmental demand is theoretically possible, as shown in Appendix A.4, but requires unphysical parameter values (either *Dλ, Dν*_*m*_ or *ν*_*m*_*ν*_*c*_ must tend to infinity). This analysis shows that perfect adaptation is not possible with reasonable parameters in biology.

**Figure 4:**
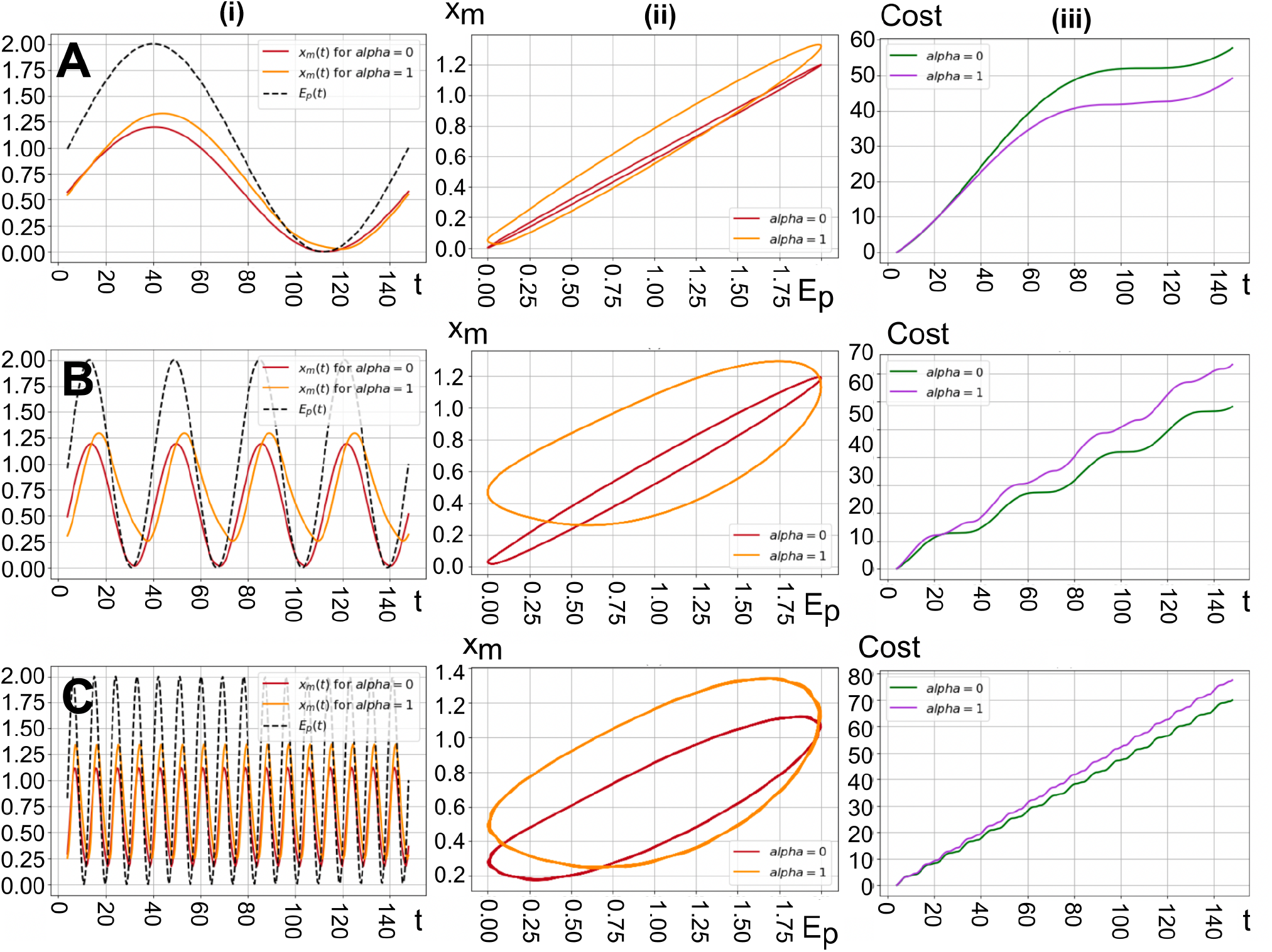
Time behaviour and phase portraits of the system in different examples of periodically changing environments. Different rows give different environmental frequencies *k* progressing vertically up Figure 3: **(A)** *k* = 1; **(B)** *k* = 4; **(C)** *k* = 16. Left column (i) shows time series of environmental demand *E*(*t*) and *x*_*m*_(*t*) for both encoding compartments. Central column (ii) shows demand vs supply traces; departure from the diagonal corresponds to a supplydemand imbalance. Right column (iii) shows time series of accumulated costs. In each case the time series are plotted after equilibration (see Methods). The common parameter values in all cases are a synthesis rate *λ* = 1, a degradation rate in the organelle *ν*_*m*_ = 0.5, no degradation in the cytosol *ν*_*c*_ = 0, a wild-type proportion of oDNA *p* = 0.75, time-averaged energetic demand *a* = 1 and relative amplitude *b* = 1. In (*i*) the transport is *D* = 0.1 and the frequency *k* = 1, in (*ii*)*D* = 0.1 and *k* = 4 and in (*iii*)*D* = 1 and *k* = 16.

### 3.4 Connection with eukaryotic taxa

In a periodically changing environment Eq (4), the frequency *k* gives the number of environmental oscillations per day: corresponding to diurnal oscillation if *k* = 1, or to semidiurnal oscillation if *k* = 2. For several plausible parameterisations of genetic properties (rates of expression, import, and degradation), these *k* values fall in the region where organelle encoding is most favoured. In particular, organelle encoding is strongly favoured in the face of diurnal or semidiurnal oscillation for genes with low *D*, that is, those that are challenging to import to the organelle (perhaps due to hydrophobicity (Björkholm et al., 2015; von Heijne, 1986)). In such cases, we expect species experiencing strong diurnal or semidiurnal variation in their organelle demands to favour organelle encoding of sets of oDNA genes that appear in nuclear DNA in other species.

To explore the feasibility of this picture, we extracted the distributions of retained gene counts across all eukaryotes with sequenced organelle DNA (see Methods). Corresponding to our original hypothesis, and the diurnal-semidiurnal predictions of this theory, we asked whether oDNA gene counts in species subjected to such environmental oscillations were higher than those in environments with different (and particularly more limited) dynamics.

Figure 5A-B show the statistics of oDNA gene counts by eukaryotic clade. First, intracellular parasites largely exist inside the highly buffered cells of their host, presented with constant, low demand on their organelles. Correspondingly, they retain very few oDNA genes. This is visible in the mtDNA counts of apicomplexans and subsets of fungi and metazoa. Apicomplexans also possess a plastid-like organelle called the apicoplast, which again contains very few oDNA genes. Free-living fungi often exist on relatively stable nutrient sources like decaying wood or leaf matter (posing few temporal fluctuations in demand), and also retain few mtDNA genes. Plants (within Viridiplantae) are typically subject to diurnal light and temperature changes and, being sessile, cannot move away from other environmental changes, and they retain comparatively many oDNA genes. Sessile, intertidal eukaryotes (including multicellular red and brown algae) face both strong diurnal light fluctuations and semidiurnal tidal variation in oxygen, temperature, and salinity, and they retain the highest plastid DNA (ptDNA) counts (Giannakis et al., 2022). There will certainly be other factors influencing where a given gene is encoded, including the evolutionary history of a species and many other features. We here just aim to draw attention to some examples which are compatible with the predictions of our theory.

**Figure 5:**
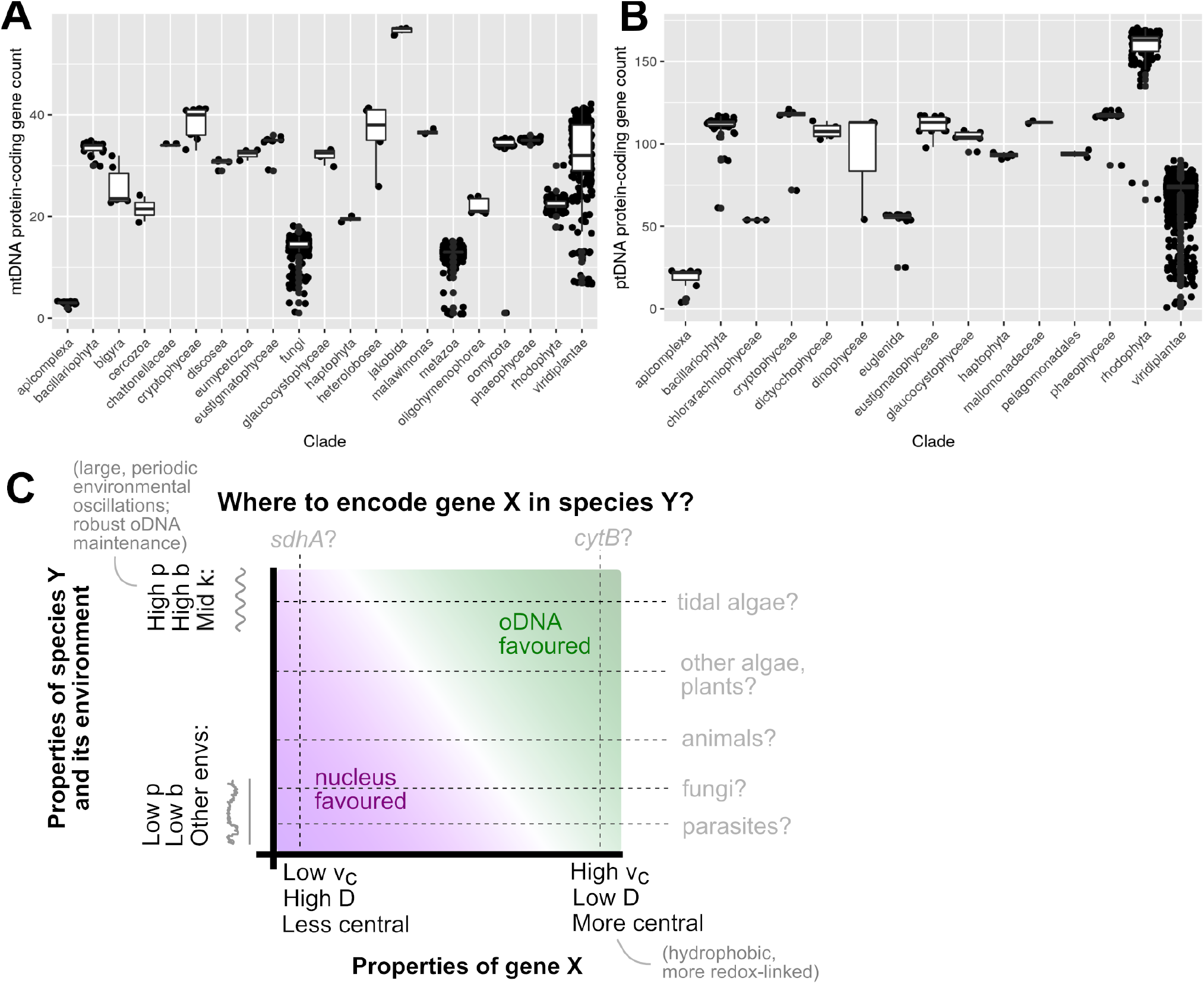
Connecting theory to oDNA profiles of extant eukaryotic taxa. **(A-B)** Protein-coding oDNA gene counts in different eukaryotic clades. (A) mtDNA, (B) ptDNA. The lower points in fungi, metazoa, and viridiplantae (for both organelles) typically correspond to parasitic species, and Apicomplexa are all parasites. **(C)** Summary illustration of the key proposed influences suggested by our model. Both cell biological and environmental features influence which compartment is preferred for encoding a given gene: the figure condenses several degrees of freedom into a single illustrative axis in both cases. At the species level, organisms maintaining high genetic integrity (high *p*; assisted, for example, by developmental bottlenecks and recombination (Edwards et al., 2021)) and subject to strong, intermediate frequency environmental oscillations (high *b*, intermediate *k*; for example, diurnal light or tidal oscillations) retain more genes. At the gene level, genes challenging to import to the organelle (high *ν*_*c*_, low *D*; for example, hydrophobic gene products) and playing a more central role in bioenergetic supply (Allen and Martin, 2016; Giannakis et al., 2022) are the more likely to be retained. Transects through the figure give examples of eukaryotic clades and mtDNA genes (generally oDNA-encoded *cytB* and generally nuclear-encoded *sdhA*) corresponding qualitatively to different cases.

Our theory suggests that the tradeoff between response to environmental demands and maintaining genetic integrity will determine which compartment is favoured for a given gene in a given organism of a specific species. In addition to the different environments described above, different species maintain oDNA integrity to different extents. Plants have lower sequence mutation rates in oDNA than nDNA (although they have pronounced structural diversity in oDNA) (Johnston, 2019a), due at least in part for their capacity for oDNA recombination (Edwards et al., 2021; Chen, 2013), suggesting that their *p* in our model would be higher and organelle encoding would be further favoured. Metazoa and fungi have high oDNA mutation rates, suggesting a lower *p* and corresponding relative favouring of nuclear encoding.

## 4 Discussion

Our model describes a link between the environmental dynamics that an organism of a given species faces and the extent to which organelle gene retention is favoured (Figure 5C). Holding other factors equal, organisms experiencing high-amplitude and/or intermediate-frequency oscillations in environmental demand will favour oDNA encoding of genes more than organisms in less periodic environments. The scale of the rates and values involved mean that this picture could help explain the high oDNA retention counts of organisms in tidal environments and subject to diurnal light oscillations, and the low retention counts in organisms in buffered, stable, or noisy environments. To our knowledge, this is the first theoretical approach attempting to explain these patterns. Of note is the connection between biophysical features governing the cell biological behaviour of gene products and the ability of the cell to respond to changing environments: a connection between the hydrophobicity (von Heijne, 1986; Björkholm et al., 2015) and CoRR (Allen and Raven, 1996; Allen, 2015) hypotheses, often viewed as competing (Giannakis et al., 2022).

A natural extension to the work presented throughout this paper is to consider oDNA damage inheritance and buildup in an evolutionary time-scale, instead of analysing the system for shorter, ecological time-scales where the proportion of damaged oDNA 1 −*p* is maintained constant. Another natural target for future work involves the transitions between encoding compartments, as any shift of an oDNA gene to the nucleus will require a transient state in which the gene is encoded in both compartments. Future theoretical work is aimed to address these questions. As with most theories about large-scale evolutionary processes, a direct connection with experimental testing is challenging. It may be possible to experimentally test these ideas further by subjecting otherwise similar organisms that differ in oDNA retention profiles to artificial fluctuating environments and assaying their performance, for example, algae in fluctuating light conditions.

We do not (and cannot) claim that this theory explains the full diversity of oDNA retention profiles across eukaryotes: as with parallel gene-specific questions, it is likely that several factors contribute together. However, although the present-day oDNA gene profile of an organism is shaped by many factors over its entire evolutionary history (including serial endosymbiosis events in the case of plastids (Keeling, 2010)), and not just its present-day situation (Janouškovec et al., 2017), some biological observations do at least qualitatively suggest that the compatibility of gene profiles of modern organisms and their environments does not conflict with our predictions (Figure 5). Recent work has shown that the transfer of mtDNA to the nucleus is ongoing in modern humans, estimating that one in every few thousand births experiences a *de novo* transfer of mtDNA material (Wei et al., 2022). It is known that transfer of ptDNA to the nucleus occurs frequently even over an individual plant’s lifetime (Stegemann et al., 2003). This ever-present potential for gene transfer suggests that modern organisms’ environments may indeed provide some selective advantage to the oDNA retention profiles they have adopted, and that ongoing selective pressures act to shape oDNA in modern eukaryotes.

In parallel with this species-specific picture, gene-specific features have been found to rank individual genes from ‘most potential to be retained’ to ‘least potential to be retained’ (Giannakis et al., 2022) (Figure 5C). Those with most retention potential typically encode the most hydrophobic products (corresponding in our model to a high degradation in the cytosol *ν*_*c*_ or slow import of gene product to the organelle *D*, and hence substantial loss in translocation to the organelle), or those most central to the assembly and production of respiratory or photosynthetic complexes (and hence most linked to supplying energetic demands) (Giannakis et al., 2022). This last feature is connected via by the CoRR hypothesis to the retention of organelle genes (Allen, 2015). Those with least retention potential, by contrast, may experience fewer barriers to translocation and be less essential for fulfilling environmental demands. Taking these pictures together, we can say that species in environments where retention is disfavoured will only retain those gene with highest retention potential, whereas species in environments where retention is more favoured will retain more of the lower-potential genes (Figure 5C). Our model thus connects cell biology, environmental demands, and genetic location to propose how specific combinations of gene and species properties shape the patterns of oDNA retention across species.

## Acknowledgments

This project has received funding from the European Research Council (ERC) under the European Union’s Horizon 2020 research and innovation programme (Grant agreement No. 805046 (EvoConBiO) to IGJ). IGJ gratefully acknowledges support from the Peder Sather Center.

## Supplementary Information

### A Study of the dynamics

In this section we carry out a standard study of the dynamics of our model Eq (2) described in Methods. We look at the well-posedness of the model in A.1, equilibrium points in A.2, stability of solutions in A.3 and we also find expressions of the solutions using different approaches for the different branches and levels of complexity of the model in A.4.

#### A.1 Well-posed model

A mathematical model is well-posed if a solution exists, the solution is unique for a set of initial conditions, and the behaviour of the solution changes continuously with respect to the initial conditions. The existence and uniqueness of solution of our model 2 for the deterministic environment *E*_*p*_(*t*) Eq (4) follows from Picard-Lindelöf theorem (Theorem I.3.1 in (Coddington and Levinson, 1955)), and it is straightforward to see that the solution changes continuously with respect to the initial conditions. Note that in Eq (4) we set the characteristic time-scale *τ* = 1 for simplicity. For the stochastic environments *E*_*w*_(*t*) Eq (5) and *E*_*r*_(*t*) Eq (6), we need other approaches that we do not include here.

**Theorem A.1** (Picard-Lindelöf theorem). *Let D* ⊆ ℝ × ℝ^*n*^ *be a closed rectangle with* (*t*_0_, *y*_0_) ∈ *D. Let f* : *D* → *ℝ*^*n*^ *be a function that is continuous in t and Lipschtiz continuous in y. Then there exists some ϵ >* 0 *such that the initial value problem y*^′^(*t*) = *f* (*t, y*(*t*)), *y*(*t*_0_) = *y*_0_ *has a unique solution y*(*t*) *on the interval* [*t*_0_ − *ϵ, t*_0_ + *ϵ*].

For the organelle-encoding case,

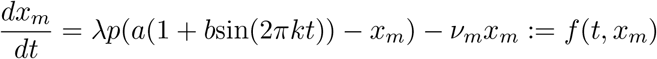

with *x*_*m*_(0) = 0, so that (0, 0) ∈ *D* ⊆ *ℝ* × ℝ. It is clear that *f* (*t, x*_*m*_) is continuous in *t* because the sine function is continuous. To check that *f* (*t, x*_*m*_) is Lipschtiz continuous in *x*_*m*_, we look if its derivative with respect to *x*_*m*_ is bounded (this implication is straightforward by the fundamental theorem of calculus):

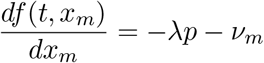

which is constant and therefore bounded.

For the nuclear-encoding case,

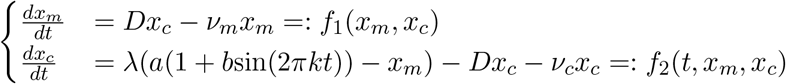

with initial conditions *x*_*m*_(0) = *x*_*c*_(0) = 0. The function *f*_1_ is continuous in *t* as it is independent of *t*, and *f*_2_ is continuous in *t* because the sine function is continuous. We check that the derivatives of *f*_1_ and *f*_2_ with respect to the state vector *y* = (*x*_*m*_, *x*_*c*_) are bounded, so that both functions are Lipschtiz continuous in *y*:

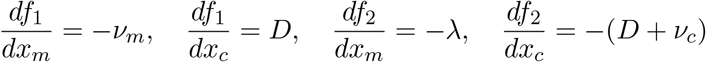

are all constant and therefore bounded.

For both cases any *ϵ >* 0 works so that [−*ϵ, ϵ*] is the interval where the unique solution is found. However, recall that *x*_*m*_, *x*_*c*_ ≥ 0, so that the solution interval is in practice restricted to [0, *ϵ*] for some *ϵ >* 0.

#### A.2 Equilibrium points

For the organelle-encoding case and a constant environment *E*(*t*) = *a* the model is

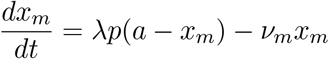

so that the equilibrium point is 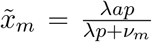. This steady state can be seen as a line whose dependent variable is *a*, so that we can write it as 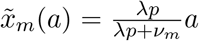 and since *ν*_*m*_ ≥ 0, we have a positive slope in the lower diagonal of [0, ∞) × [0, ∞).

For the nuclear-encoding case and a constant environment *E*(*t*) = *a* the model is

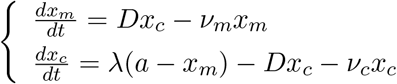

with equilibrium point

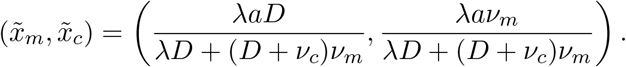

#### A.3 Stability analysis

We set the environment constant *E*(*t*) = *a* for simplicity, but the same analysis holds for a fluctuating environment *E*(*t*) = *E*_*p*_(*t*) as defined in Eq (4). For the organelle-encoding case and the branch of the model *f* (*E, x*_*m*_) = *a* − *x*_*m*_, we look at the Jacobian *J* of *λp*(*a* − *x*_*m*_) − *ν*_*m*_*x*_*m*_, which is

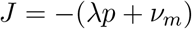

so that the unique eigenvalue is *γ* = − (*λp* + *ν*_*m*_) *<* 0, meaning that the solution is a stable node which is Lyapunov asymptotically and uniformly stable.

For the branch *f* (*E, x*_*m*_) = 0, the Jacobian of −*ν*_*m*_*x*_*m*_ is *J* = −*ν*_*m*_, so if *ν*_*m*_ ≠ 0 we have an eigenvalue *γ* = −*ν*_*m*_ *<* 0, meaning that the solution is a stable node which is Lyapunov asymptotically and uniformly stable. If *ν*_*m*_ = 0, we cannot say more without further analyses.

For the nuclear-encoding case and the branch *f* (*E, x*_*m*_) = *a* − *x*_*m*_, we look at the Jacobian of

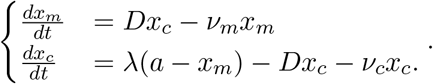

which is

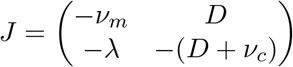

so that the eigenvalues are 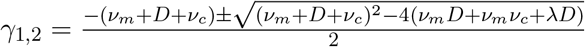

Here we have to study the sign of the real part of *γ*_1,2_. If the interior of the square root is 0, then we have a double negative real eigenvalue and a degeneracy which requires further analysis. This is the case when 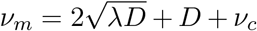. Note that this case does not hold for the simulations in Figure 3 as for fixed *ν*_*m*_ = 0.5 and *ν*_*c*_ = 0, *D* must be equal to 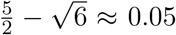, but *D* is set to take orders of magnitudes 10^*i*^ for *i* = −2, −1, 0.

If the interior of the square root is negative, then we have two distinct complex eigenvalues whose real part is negative as long as one of the parameters *ν*_*m*_, *ν*_*c*_ or *D* is greater than zero (which is the case of *ν*_*m*_ and *D* in the simulations in Figure 3), and we have a stable spiral which is Lyapunov asymptotically and uniformly stable. This is the case if *λ, D >* 0 and 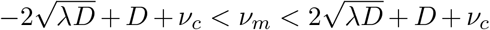. In the simulations in Figure 3 where *λ* = 1, *ν*_*m*_ = 0.5 and *ν*_*c*_ = 0, we see that these inequalities hold for *D* = 0.1 and *D* = 1, but fail for *D* = 0.01. For the standard case of setting all parameters equal to 1, the model also falls into this classification of stable spiral, as the eigenvalues are 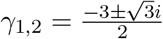.

If the interior of the square root is positive, then we might have a case of instability if there are two distinct real eigenvalues where one is negative and the other one is positive.

The interior of the square root is positive if *λ >* 0 and either (*i*) 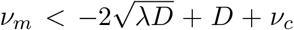 or (*ii*) 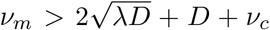. Since 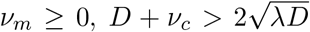 so that (*i*) can hold. This is the case if *D* is low, *λ* is low and *ν*_*c*_ is high, for example for *D* = 0.01, *λ* = 0.01 and *ν*_*c*_ = 1, *ν*_*m*_ *<* 0.98 and since in the simulations *ν*_*m*_ ∈ [0, 1], this covers most of the parameter values of *ν*_*m*_. Case (*ii*) cannot hold for *ν*_*m*_ = 0. However, case (*ii*) holds for the simulations in Figure 3 for *D* = 0.01, *λ* = 1, *ν*_*m*_ = 0.5 and *ν*_*c*_ = 0. We plug these values in *γ*_1,2_ and see that there are two distinct negative real eigenvalues, so the solution is a stable node which is Lyapunov asymptotically and uniformly stable.

Lastly, if *ν*_*m*_ = *D* = *ν*_*c*_ = 0, then *γ*_1,2_ = 0 and we cannot say more about the stability of the model. For the branch *f* (*E, x*_*m*_) = 0, the Jacobian of

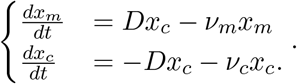

is

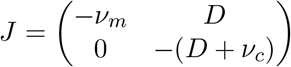

so that we have two eigenvalues *γ*_1_ = −*ν*_*m*_ *<* 0 and *γ*_2_ = − (*D* + *ν*_*c*_) *<* 0, and the solution is therefore a stable node Lyapunov asymptotically and uniformly stable.

The idea of a solution being Lyapunov uniformly stable is that it stays close to another solution for a different set of initial conditions, and of being Lyapunov asymptotically stable is that it stays close in time to another solution for a different set of initial conditions, so that the distance between both solutions converge to 0 when *t* tends to infinity.

This study of the stability of solutions shows that our biological system is stable. In particular, the variables *x*_*m*_(*t*) and *x*_*c*_(*t*) representing the amount of gene product in the organelle and in the cytosol, respectively, are bounded below by 0 to have such a meaning in biology, but this stability analysis is saying that *x*_*m*_(*t*) and *x*_*c*_(*t*) are also bounded above. Therefore, the system does not explode when *t* tends to infinity.

#### A.4 Expressions of solutions

The model of ODEs presented in Eq (2) for *E*(*t*) = *E*_*p*_(*t*) defined in Eq (4) is

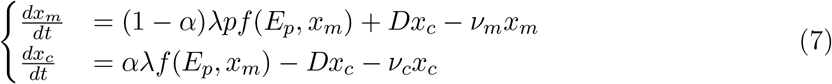

The model has two branches, one for *f* (*E*_*p*_(*t*), *x*_*m*_(*t*)) = *E*_*p*_(*t*) −*x*_*m*_(*t*) and another for *f* (*E*_*p*_(*t*), *x*_*m*_(*t*)) = 0. For this last one, the system enters in decay, and we can solve it in a straightforward way as follows.

For the organelle-encoding case (*α* = 0) we have the ODE

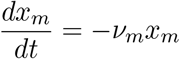

which has solution

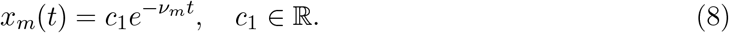

For the nuclear-encoding case (*α* = 1) we have the ODEs

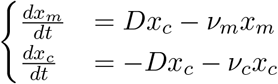

with solutions

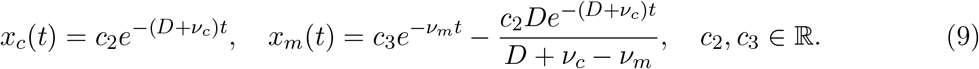

For the other branch *f* (*E*_*p*_(*t*), *x*_*m*_(*t*)) = *E*_*p*_(*t*) −*x*_*m*_(*t*), we first look when the environment is constant equal to *a*. For the organelle-encoding case we get the solution

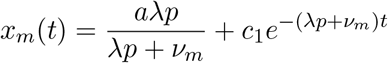

and since *x*_*m*_(0) = 0,

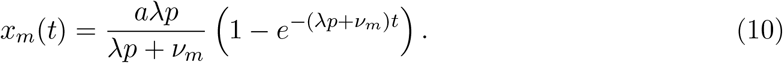

For the nuclear-encoding case and a constant environment, the solutions look too long and not so informative to be presented here.

Next we look at this same branch *f* (*E*_*p*_(*t*), *x*_*m*_(*t*)) = *E*_*p*_(*t*) − *x*_*m*_(*t*) for a periodically changing environment. This case is very hard to compute, so we use a different approach consisting on finding coefficients of sinusoidal waves that give the expressions of the solutions of the variables *x*_*m*_(*t*) and *x*_*c*_(*t*) (as we see in the simulations in Figure 4 this sinusoidal form that the solutions have). Moreover, we compare these solutions to the environmental wave to see for which parameter values the system is in-phase or out-of-phase. This approach is divided in the next two subsections, first determining the coefficients of the sinusoidal solutions and then comparing them to the periodic environmental wave *E*_*p*_(*t*).

##### A.4.1 Determining coefficients of solutions

For a periodically changing environment *E*_*p*_(*t*) defined in Eq (4) with *τ* = 1 and *b* ≠ 0

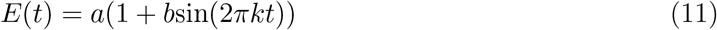

we want to find expressions of solutions for the branch of the model corresponding to *f* (*E*_*p*_(*t*), *x*_*m*_(*t*)) = *E*_*p*_(*t*) −*x*_*m*_(*t*). For simplicity, we set *p* = 1 in the ODEs Eq (2). From the simulations in Figure 4 we observe that the variables follow a sinusoidal wave in time, so we want to determine the coefficients of a sum of sine and cosine functions that capture the dynamics of the solutions.

For the organelle-encoding case (*α* = 0), we write *x*_*m*_(*t*) as

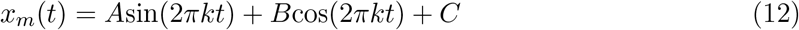

where *A, B, C* are the constants to determine.

From the system of ODEs Eq (2) we have that the derivative of *x*_*m*_(*t*) is

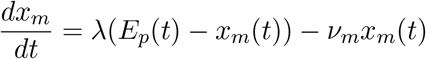

and by substituting by Eqs (12) and (11) we have

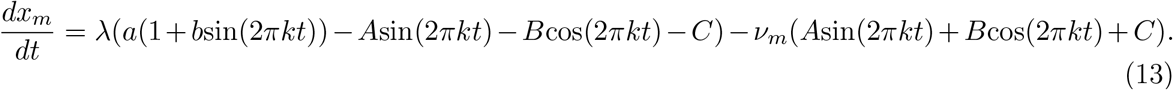

We then derivate Eq (12) and we set it equal to the right-hand side of Eq (13), getting the constants

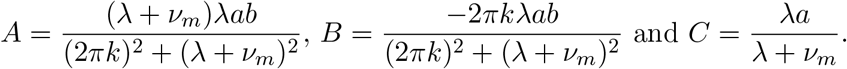

Hence

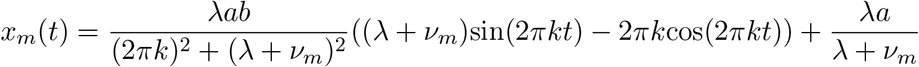

and since 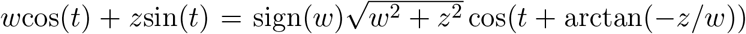, the equation above becomes

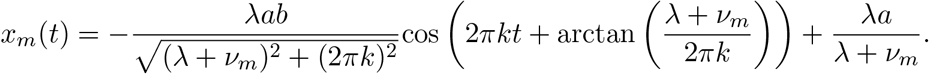

To compare more easily the variable *x*_*m*_(*t*) with the environmental function *E*_*p*_(*t*), we write *x*_*m*_(*t*) as a sine function, as it is known that cos(*t*) = sin(*t* + *π/*2) and sin(*t*) = sin(*t* + *π*), so that

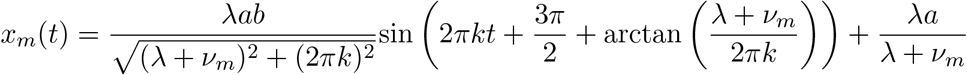

and by rearranging we get the following expression in the same form as *E*_*p*_(*t*) in Equation (11):

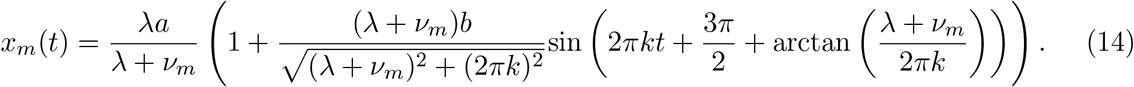

For the nuclear-encoding case (*α* = 1), the solution of the system of ODEs Eq (2) is of the form

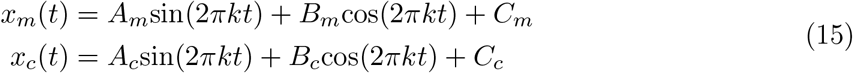

where *A*_*i*_, *B*_*i*_, *C*_*i*_ are the constants to determine, for *i* = *m, c*.

Proceeding as for the organelle-encoding case, we get

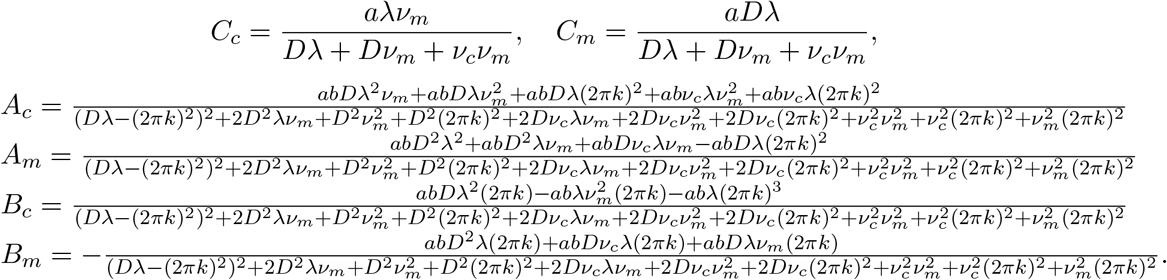

Note that *C*_*c*_, *C*_*m*_, *A*_*c*_ *>* 0, *B*_*m*_ *<* 0 and *A*_*m*_, *B*_*c*_ can change sign.

Using the expressions 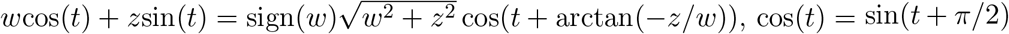 and −sin(*t*) = sin(*t* + *π*), we get a solution for *x*_*m*_(*t*) and two possible solutions for *x*_*c*_(*t*) of the form:

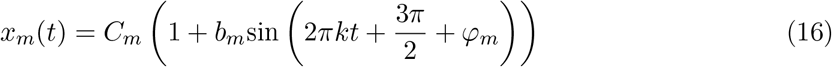

Where 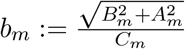and

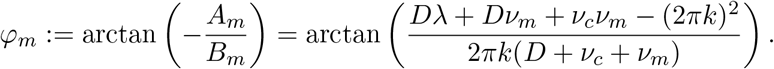

1. If 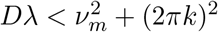 so that *B*_*c*_ *<* 0, then

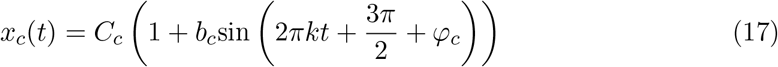

where 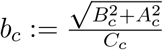 and

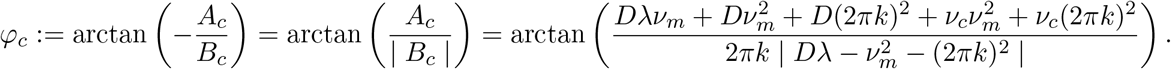
2. If 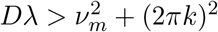so that *B*_*c*_ *>* 0, then

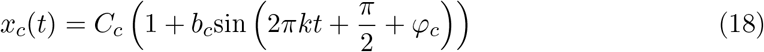

where 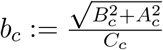 and

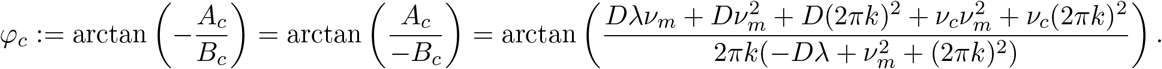

##### A.4.2 Determining the synchronicity of the organism with the environment

After finding the constants of the expressions for *x*_*m*_(*t*) and *x*_*c*_(*t*), we proceed studying when these expressions in the form of waves coincide with the environmental wave *E*_*p*_(*t*).

We look at the following three aspects of *x*_*m*_(*t*), *x*_*c*_(*t*) and *E*_*p*_(*t*), to determine whether the two first solutions coincide with the environmental function or not:

1. The time-averaged demands, also called absolute magnitudes.
2. The argument of the sine functions, so that the functions are in phase or there is a phase shift making them being less or more out of phase.
3. The amplitudes.

For the organelle-encoding case (*α* = 0), the variable *x*_*m*_(*t*) is as shown in Eq (14) and the environmental function is as in Eq (11).

So for point 1. the absolute magnitudes *a* in *E*_*p*_(*t*) and 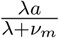 in *x*_*m*_(*t*) coincide if and only if *ν*_*m*_ = 0 and *λ* ≠ 0.

For point 2., we look at the phase shift 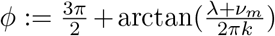. The arcotangent function in general gives values in 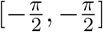, but since the argument 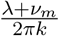 is always greater or equal than zero, arctan 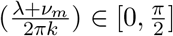. Hence,

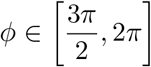

Where 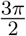 is the biggest possible phase shift and 2*π* means that there is no phase shift.

The system is hence in phase if *ϕ* = 2*π*. This is the case when *λ*+*ν*_*m*_ ∞, as arctan 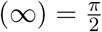, so at least one of the parameters *λ* or *ν*_*m*_ must be very large. Also, the system is in phase if we have a very low frequency *k* → 0, since 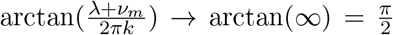. The system is completely out of phase if 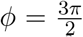, and this is the case when *λ, ν*_*m*_ → 0, so that arctan(0) = 0. We also have a phase shift 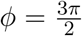 if the frequency is very high, i.e, *k* → ∞ and at least one of the parameters *λ, ν*_*m*_ is non-zero and small, or both are non-zero and small.

For point 3., the relative amplitudes *b* in *E*_*p*_(*t*) and 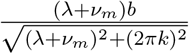 in *x*_*m*_(*t*) coincide if and only if 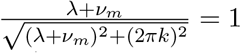, and this is the case when *k* = 0 or *k* → 0, i.e, when the environment is (almost) constant.

In conclusion, *x*_*m*_(*t*) is identical to *E*_*p*_(*t*) if *ν*_*m*_ = 0 (matching absolute magnitudes), *λ* → ∞ (matching phases) and *k* = 0 or *k* → 0 (matching amplitudes). On the other side, *x*_*m*_(*t*) is different to *E*(*t*) when the gene is located in organelle if *ν*_*m*_ *>>* 0 (the absolute magnitude of *x*_*m*_(*t*) is lower than the absolute magnitude of *E*(*t*), where *λ* 0). If that holds, then a phase shift of 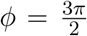 cannot hold as *λ* + *ν*_*m*_ → 0, so *k* → ∞ must be the case. The amplitude of *x*_*m*_(*t*) is also smaller than of *E*_*p*_(*t*). If *λ* + *ν*_*m*_ → 0 so that there is a phase shift of 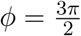, then if *ν*_*m*_ = 0 and *λ >* 0 but small, the absolute magnitudes match. The frequency must be *k*→ ∞ for the amplitudes to not coincide.

Note that we have determined that the phases and amplitudes cannot be simultaneously the same in *E*_*p*_(*t*) and *x*_*m*_(*t*) when *E*_*p*_(*t*) is not constant. This is because for being in phase both functions need to oscillate, which is the case if *λ* + *ν*_*m*_ → ∞ and *k >* 0, but for additionally having the same amplitudes, the frequency *k* must tend to 0, which contradicts the existence of a higher frequency *k >* 0 for matching phases.

For the nuclear-encoding case (*α* = 1), we compare the variable *x*_*m*_(*t*) as shown in Equation (16) with the environmental function *E*_*p*_(*t*) in Equation (11).

For point 1. the absolute magnitudes *a* in *E*_*p*_(*t*) and *C*_*m*_ in *x*_*m*_(*t*) are identical if and only if *ν*_*m*_ = 0 and *λ, D >* 0.

For point 2., we look at the phase shift

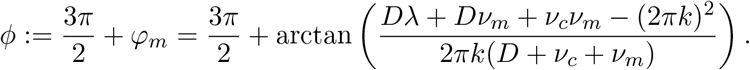

We see that for this case, *D, ν*_*m*_, *ν*_*c*_ must not all be equal to zero, and the arcotangent function gives values in 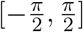, so that

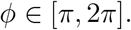

The variable *x*_*m*_(*t*) and the environmental function *E*_*p*_(*t*) are in phase when *ϕ* = 2*π*, and this holds when

- *k* → 0 or
- 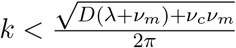 and either
  – *D* and *λ* are very large, so that *Dλ* → ∞, or
  – *D* and *ν*_*m*_ are very large, so that *Dν*_*m*_ → ∞, or
  – *ν*_*c*_ and *ν*_*m*_ are very large, so that *ν*_*c*_*ν*_*m*_ → ∞

where for each case, the rest of the parameters are not so large.

There is a phase shift of *ϕ* = *π* if *k* → ∞ and the rest of the parameters are significantly smaller than *k*.

We also see that a phase shift 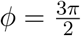is achieved when 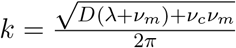

For point 3. we look when *b* = *b*_*m*_. This turns out to be very hard to solve using symbolic solvers on Python, even setting some parameters equal to 0. Therefore we do not further explore this point.

#### A.5 Nonexistence of a limit cycle

Since in the simulations in Figure 4 we see persistent oscillations in time as the solutions of our model, and closed orbits in the phase planes, it is natural to ask whether the system has a limit cycle or just a closed orbit that is not isolated. A limit cycle is defined as a closed and isolated orbit in the phase plane that corresponds to persistent oscillations in time. A periodic solution 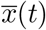 is defined such that 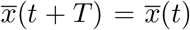 for all *t* and for some positive constant *T* called the period.

By applying Bendixon’s negative criterion, also named Bendixson–Dulac theorem, (page 318 in Burton (2005)) we determine that there are no limit cycles, just closed orbits that are not isolated from other solutions for other sets of initial conditions.

For the organelle-encoding case, we look at the general case where the proportion of nonmutant organelle DNA (oDNA) *p* has the form of an exponential decay with rate *µ*, so that this analysis works for long, evolutionary time-scales. Therefore *dp/dt* = −*µp* and we have the following system:

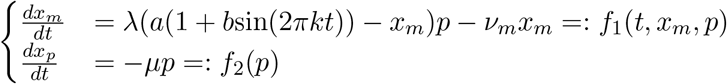

so that

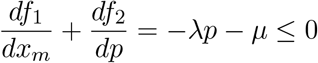

since *p* is an exponential decay with initial value *p*(0) = 1 that tends to zero, so it is always positive, and the parameters *λ, µ* are non-negative. Therefore since this expression does not change sign, by Bendixon’s negative criterion it does not exist a limit cycle.

For the nuclear-encoding case,

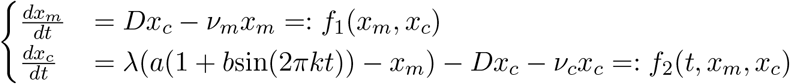

we have that

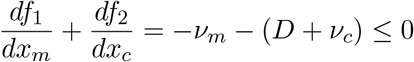

so since the expression above does not change sign, it is always less or equal than 0 since the variables are non-negative, by Bendixon’s negative criterion it does not exist a limit cycle.

## B Supplementary Figures

**Figure S1:**
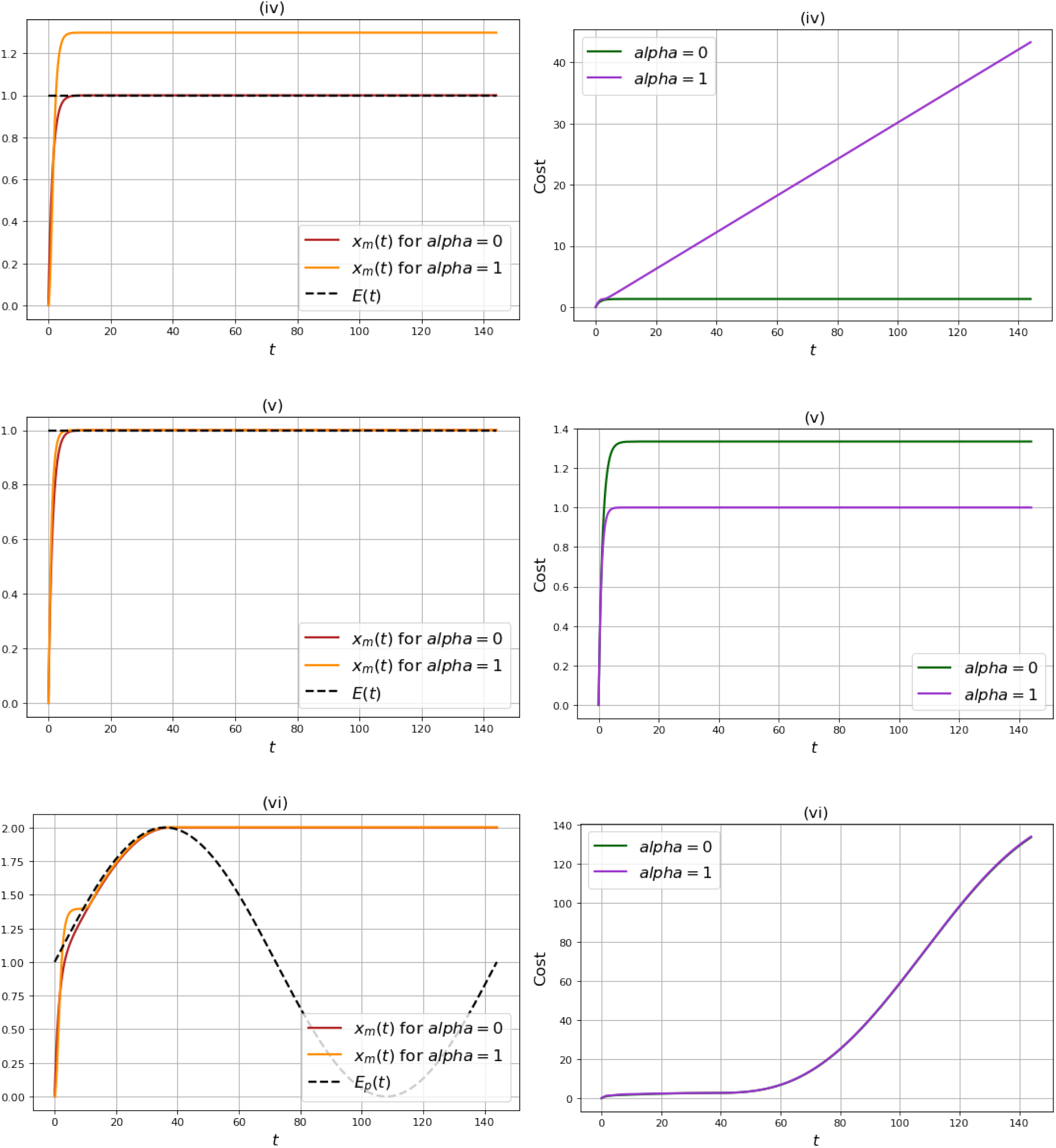
Time behaviour of the system showing the influence of the degradation rate in the organelle. In (*iv*) we see that when the gene is encoded in the organelle and the environment is constant, then if there is no degradation in the organelle (*ν*_*m*_ = 0) the system fully matches the demand after a short transient period. Other parameter values are synthesis rate *λ* = 1, transport rate *D* = 1, no degradation in the cytosol *ν*_*c*_ = 0, oDNA wild-type proprotion *p* = 0.75 and *E*(*t*) = *a* = 1. In (*v*) we get the full match between *x*_*m*_ and *E* for the nuclear-encoding case *α* = 1 by setting a faster transport *D* = 10. The rest of the parameters are as for (*iv*). In (*vi*) we have a fluctuating environment and *ν*_*m*_ = 0 to see that after fully reaching the demand, the system cannot enter into decay phase along the decreasing environmental wave, becoming out-of-phase and leading to a pronounced increase in the cost. The parameter values for this case are as in (*iv*) with *b* = 1 and *k* = 1 in *E*_*p*_(*t*) Eq(4).

**Figure S2:**
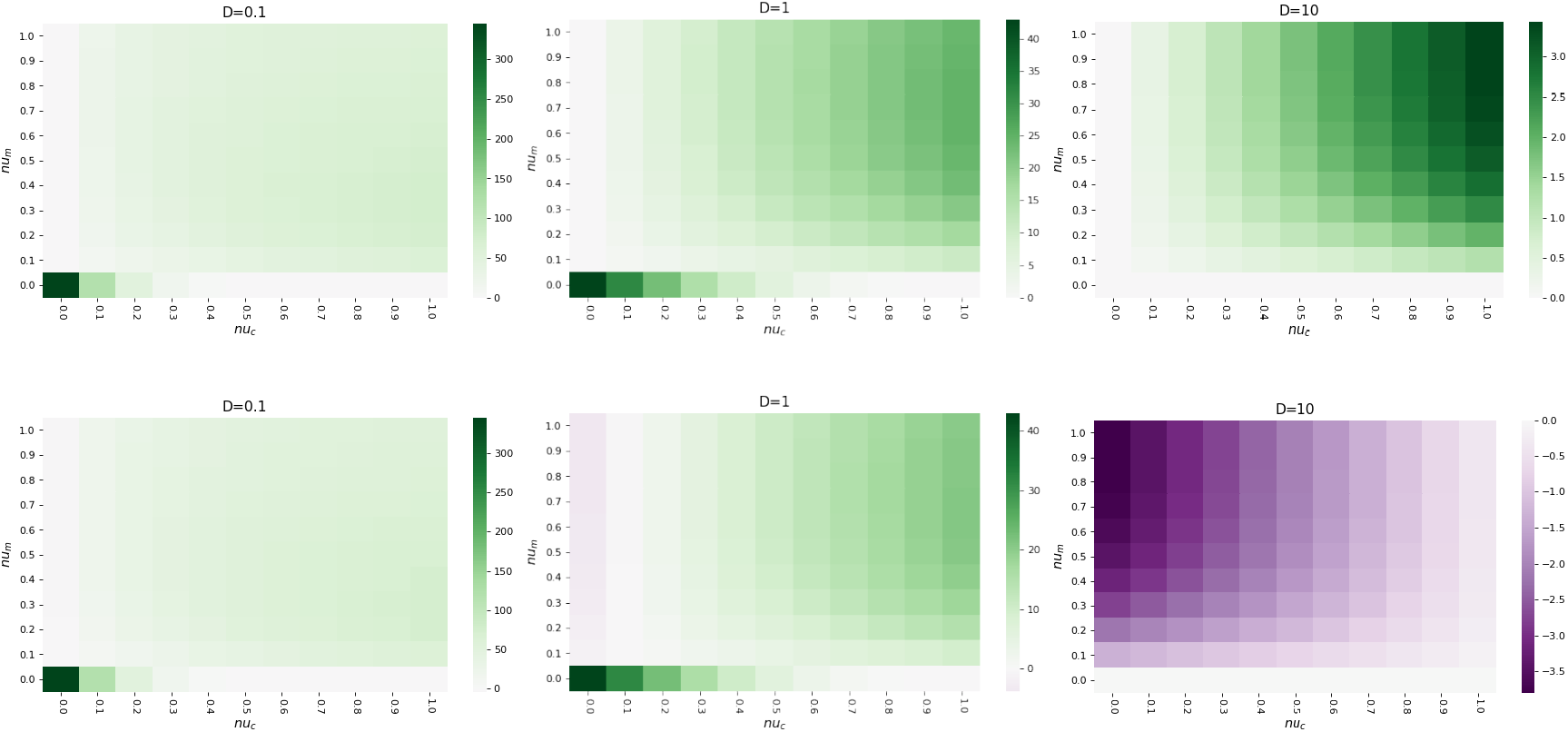
Less oDNA damage simply shifts the favoured compartment towards organelle encoding in static environments. As Figure 2, absolute cost difference in encoding strategies in static environments with cytosol degradation rate *ν*_*c*_ and organelle degradation rate *ν*_*m*_, for (top) *p* = 1; (bottom) *p* = 0.9.

**Figure S3:**
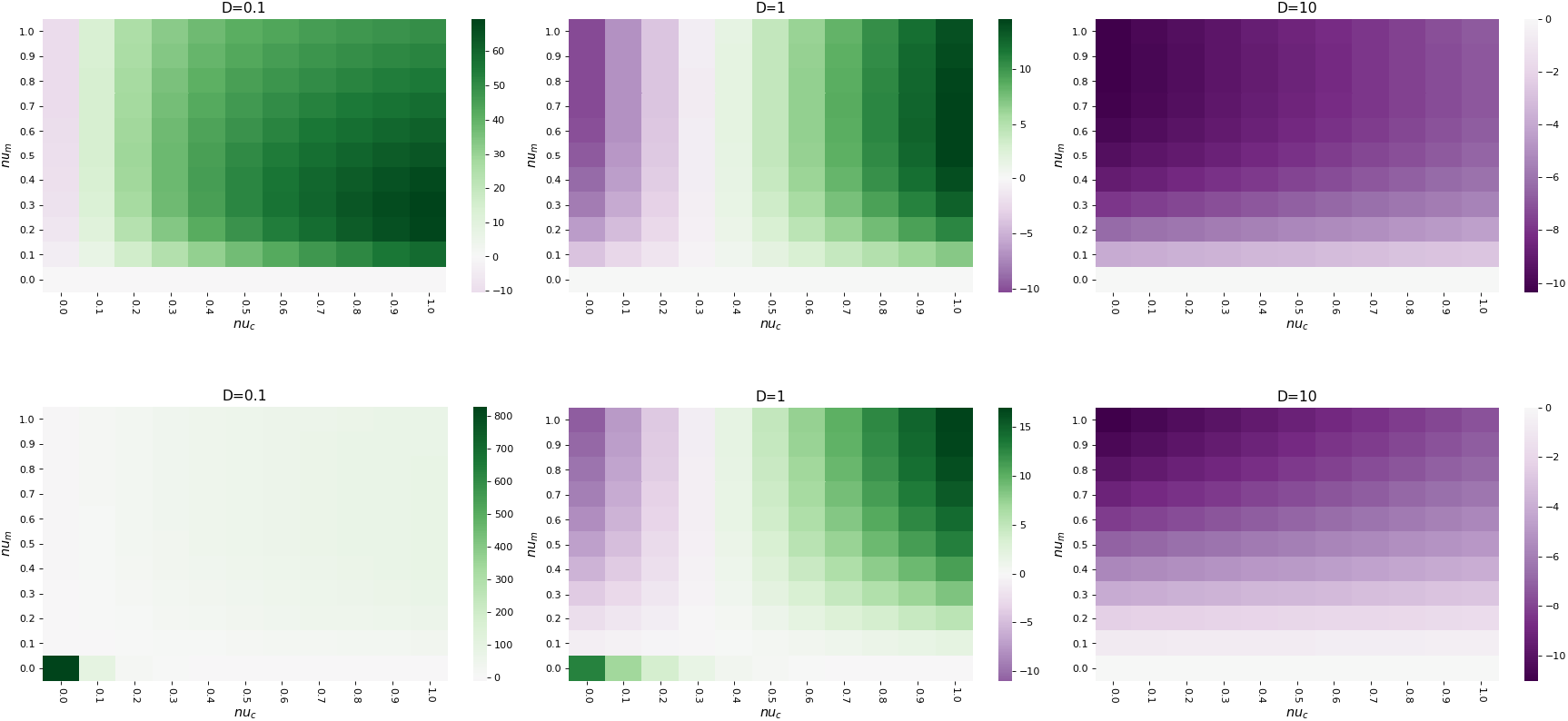
Static environments for different cost functions. As top row of Figure 2, absolute cost difference in encoding strategies in static environments with cytosol degradation rate *ν*_*c*_ and organelle degradation rate *ν*_*m*_. Top row here is for a cost defined as the integral over a time interval of the signalling function Eq (1), i.e. 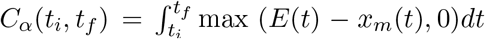. Bottom row for a cost as the signalling function Eq (1) to the square, i.e. 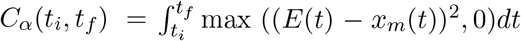. In this last case, the signalling function Eq(1) is changed to be to the square, i.e. *f* (*E*(*t*), *x*_*m*_(*t*)) = max ((*E*(*t*) −*x*_*m*_(*t*))^2^, 0). We observe that there is an outlier for *ν*_*m*_ = 0 (particularly strong when *ν*_*c*_ is also 0) when the cost is calculated with the absolute difference and the quadratic signalling function. Such an outlier does not show up when the cost is calculated with the signalling function. This is because for *ν*_*m*_ = 0, the variable *x*_*m*_(*t*) for the nuclear encoding case is significantly over the demand *E*(*t*) when *D* is low (as seen in Appendix A.2). However, with the signalling function, that difference *E*(*t*) − *x*_*m*_(*t*) is negative, so the value 0 is given, showing uniform trends without outliers across the parameter values.

**Figure S4:**
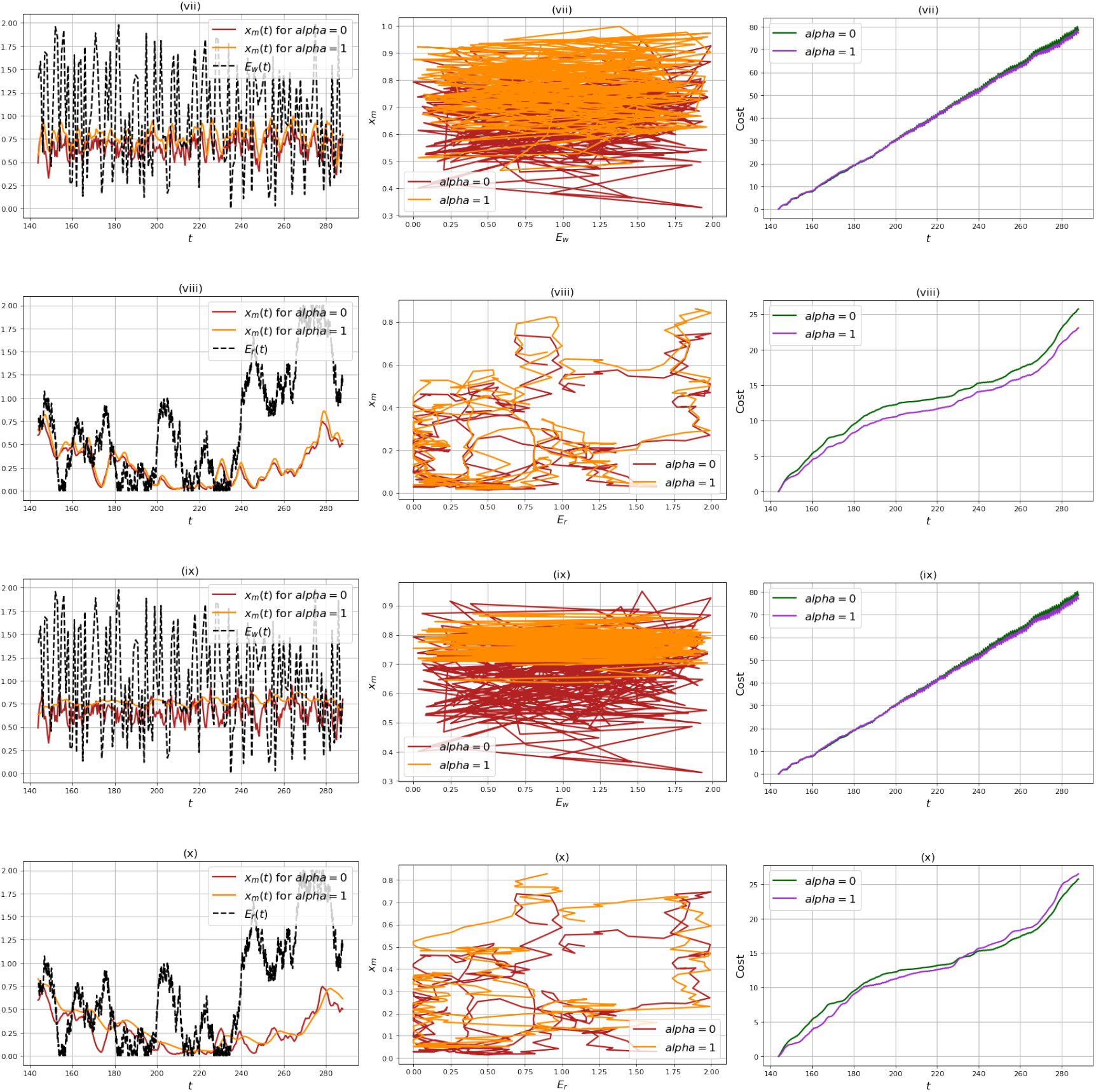
Time behaviour and phase portraits of the system in different examples of randomly changing environments. The common parameters for all three cases are a synthesis rate *λ* = 1, degradation in the organelle *ν*_*m*_ = 0.5, no degradation in the cytosol *ν*_*c*_ = 0 and wild-type proportion of oDNA *p* = 0.75. For a transport rate *D* = 1 we have white noise (*vii*) and red noise (*viii*), and for a transport rate *D* = 0.1 we have white noise (*ix*) and red noise (*x*). The left column displays the time-series and the middle columnd the phase portraits for a time window of a full day (24 hours) in time units *T* = 10min with the transient periods removed, as explained in Methods. The right column displays the accumulated costs Eq (3) in that time window.

**Figure S5:**
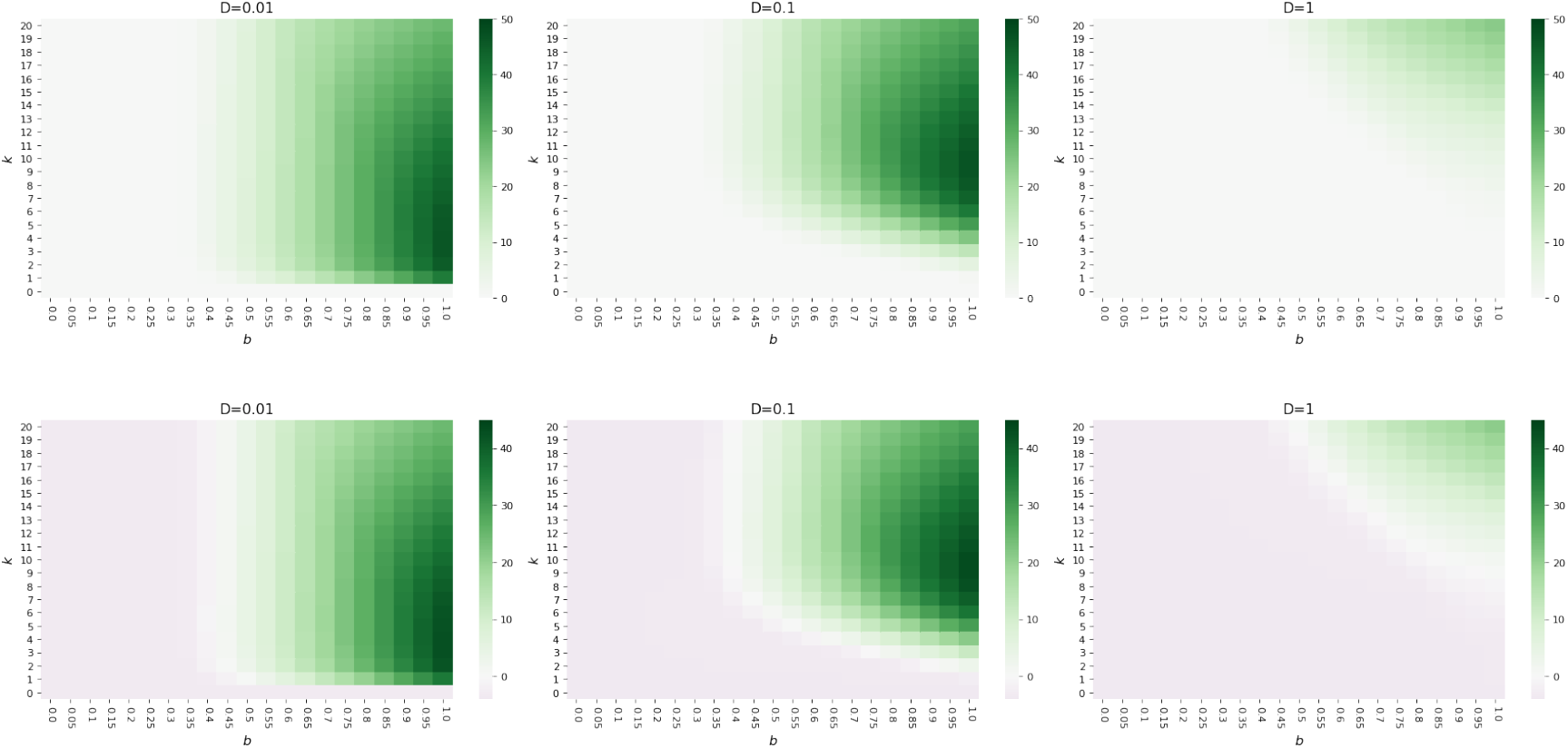
Less oDNA damage simply shifts the favoured compartment towards organelle encoding in fluctuating environments. As the bottom row of Figure 3, absolute cost difference between encoding compartments for environmental demand with oscillation amplitude *b* and frequency *k*, for (top) *p* = 1; (bottom) *p* = 0.9.

**Figure S6:**
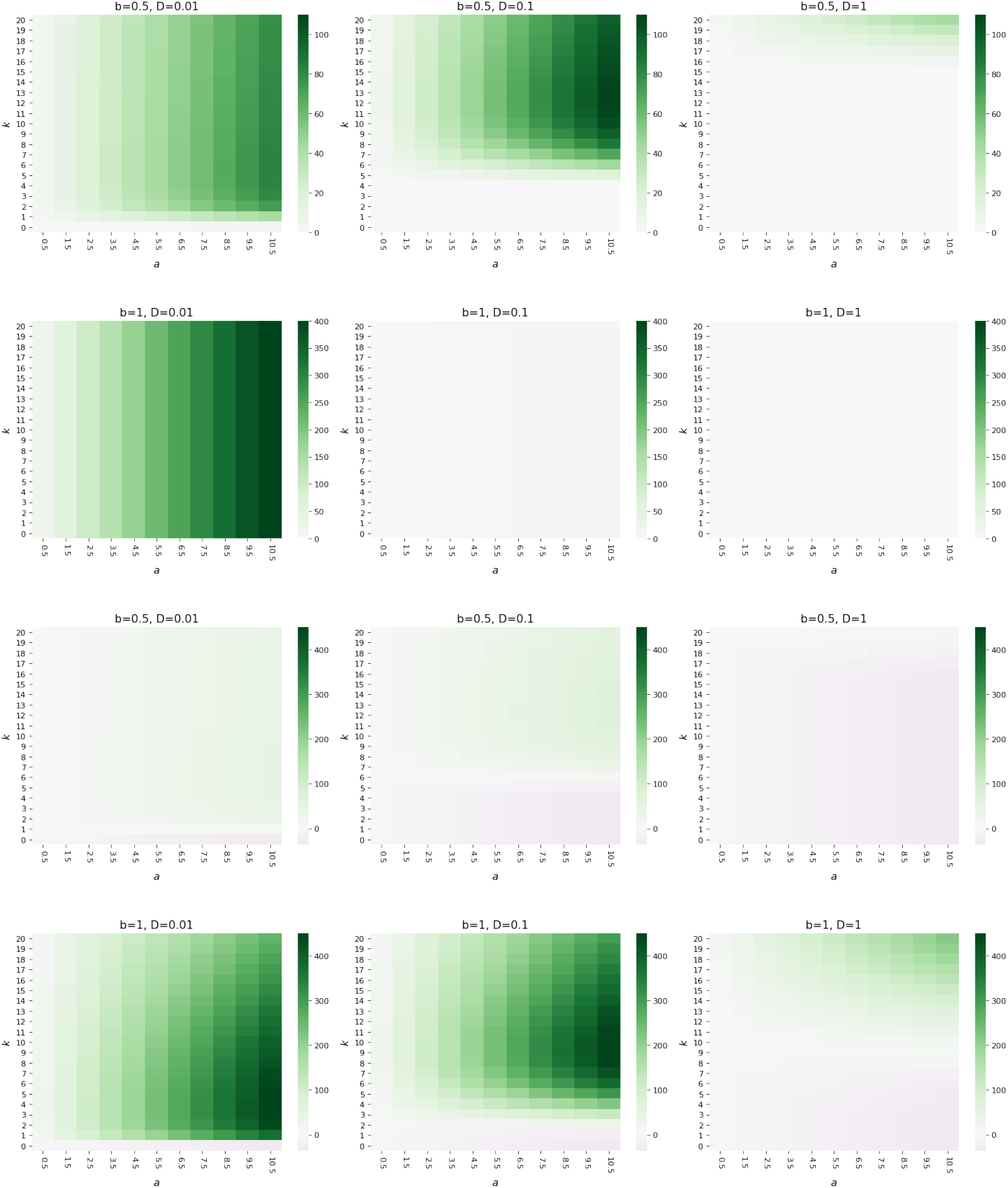
Less oDNA damage simply shifts the favoured compartment towards organelle encoding in fluctuating environments. As the top rows of Figure 3, absolute cost difference between encoding compartments for environmental demand with mean amplitude *a* and frequency *k*, for (top two rows) *p* = 1; (bottom two rows) *p* = 0.09.

**Figure S7:**
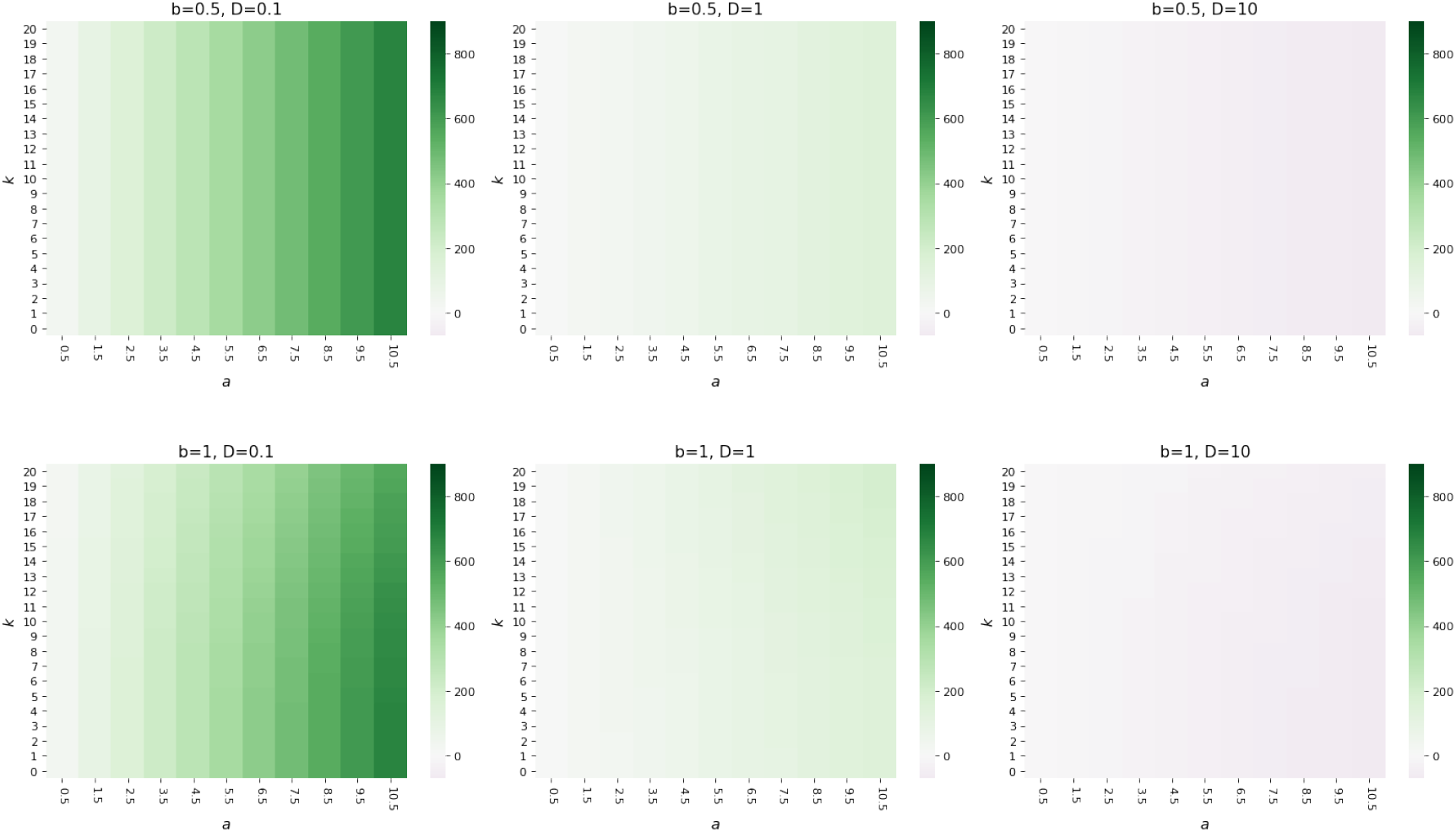
Cytoplasmic degradation simply shifts preferred compartment to organelle encoding in fluctuating environments. As the top rows of Figure 3, absolute cost difference between encoding compartments for environmental demand with mean amplitude *a* and frequency *k*, for degradation in the cytosol *ν*_*c*_ = 1. No differences for medium amplitudes *b* = 0.5 and higher amplitudes *b* = 1.

**Figure S8:**
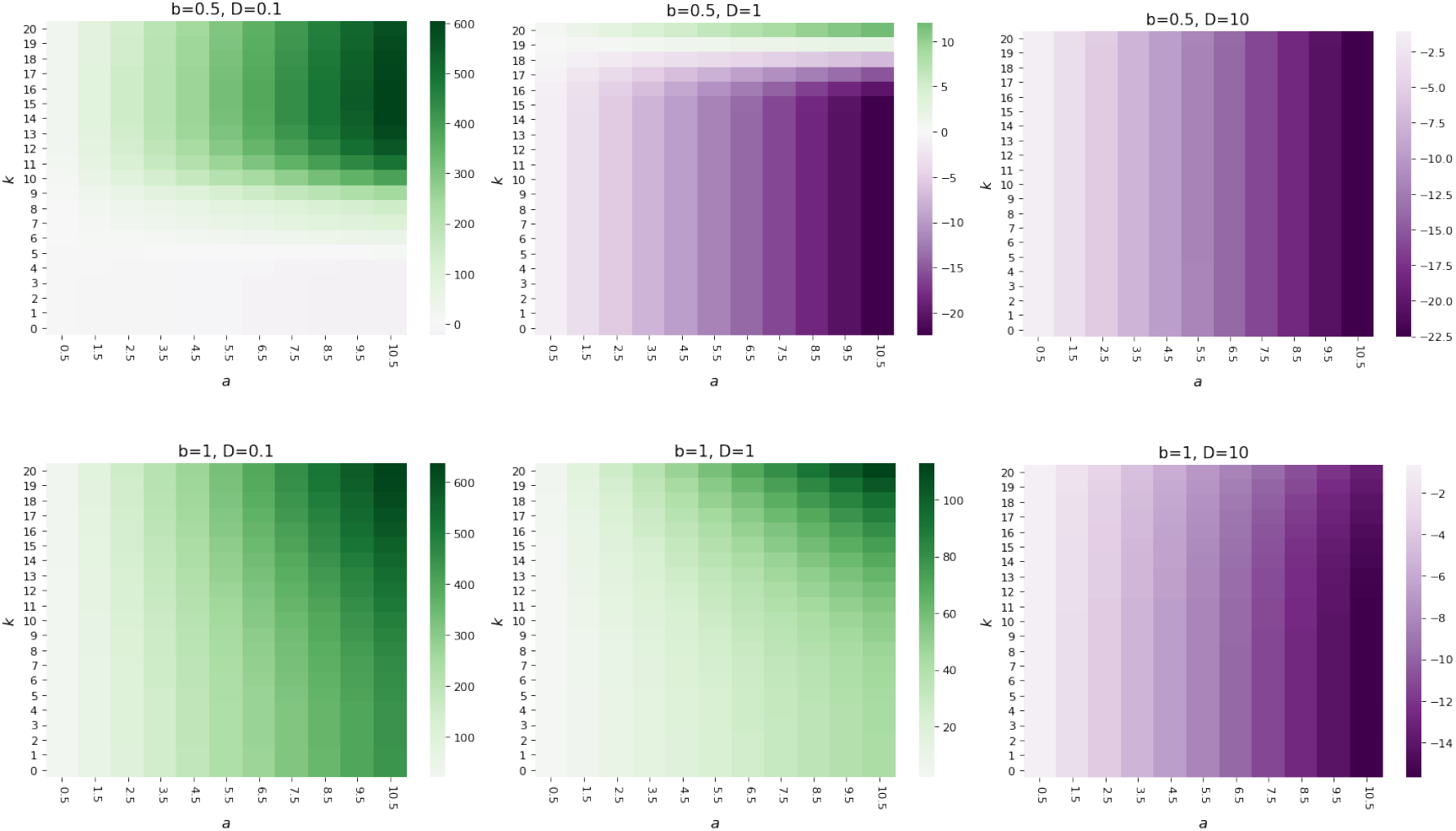
Higher synthesis rate and no cytoplasmic degradation shifts preferred compartment to organelle encoding in fluctuating environments. As the top rows of Figure 3, absolute cost difference between encoding compartments for a synthesis rate *λ* = 10.

**Figure S9:**
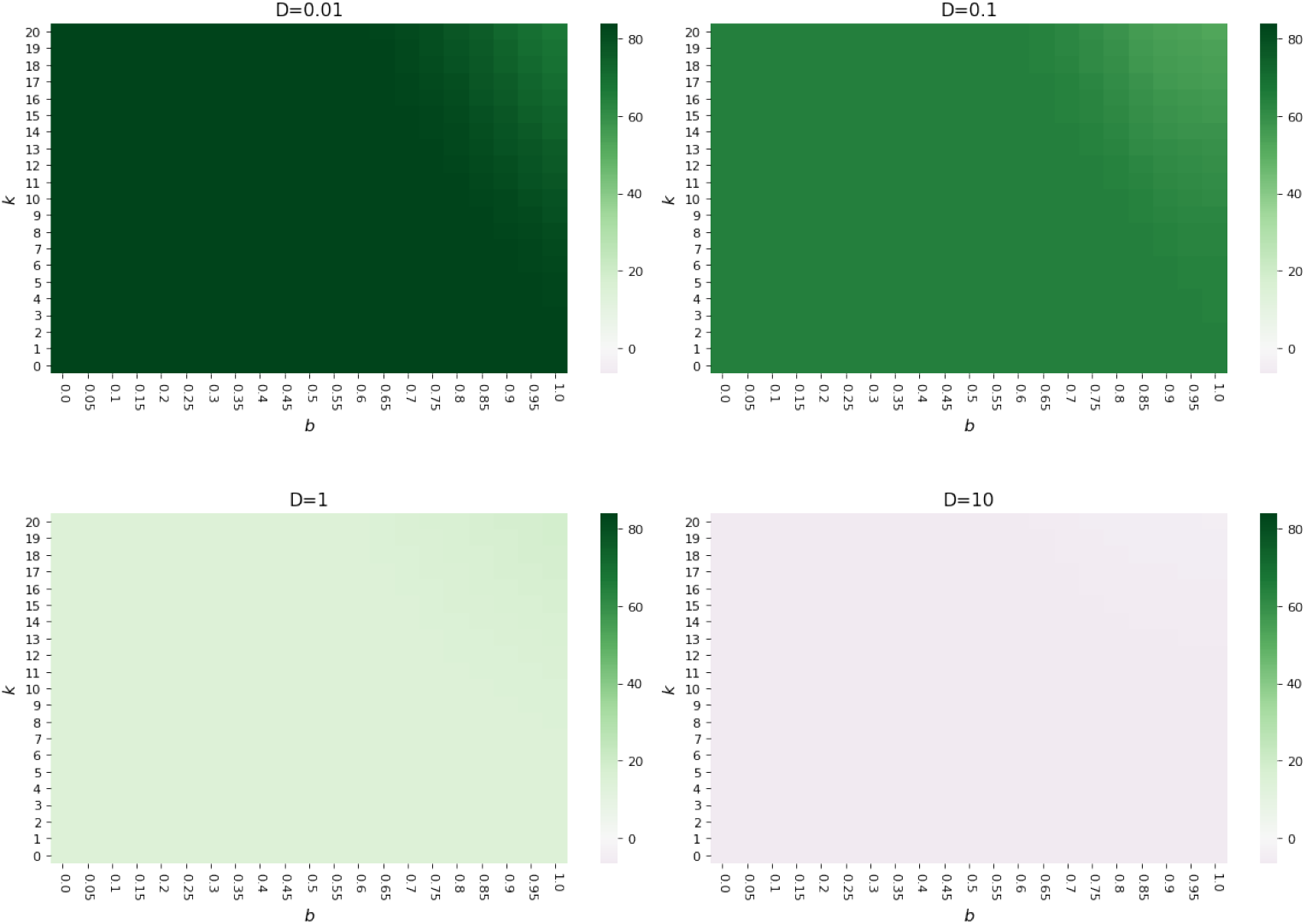
Cytoplasmic degradation simply shifts preferred compartment to organelle encoding in fluctuating environments. As the bottom row of Figure 3, absolute cost difference between encoding compartments for environmental demand with oscillation amplitude *b* and frequency *k*, for *p* = 0.75 and *ν*_*c*_ = 1.

**Figure S10:**
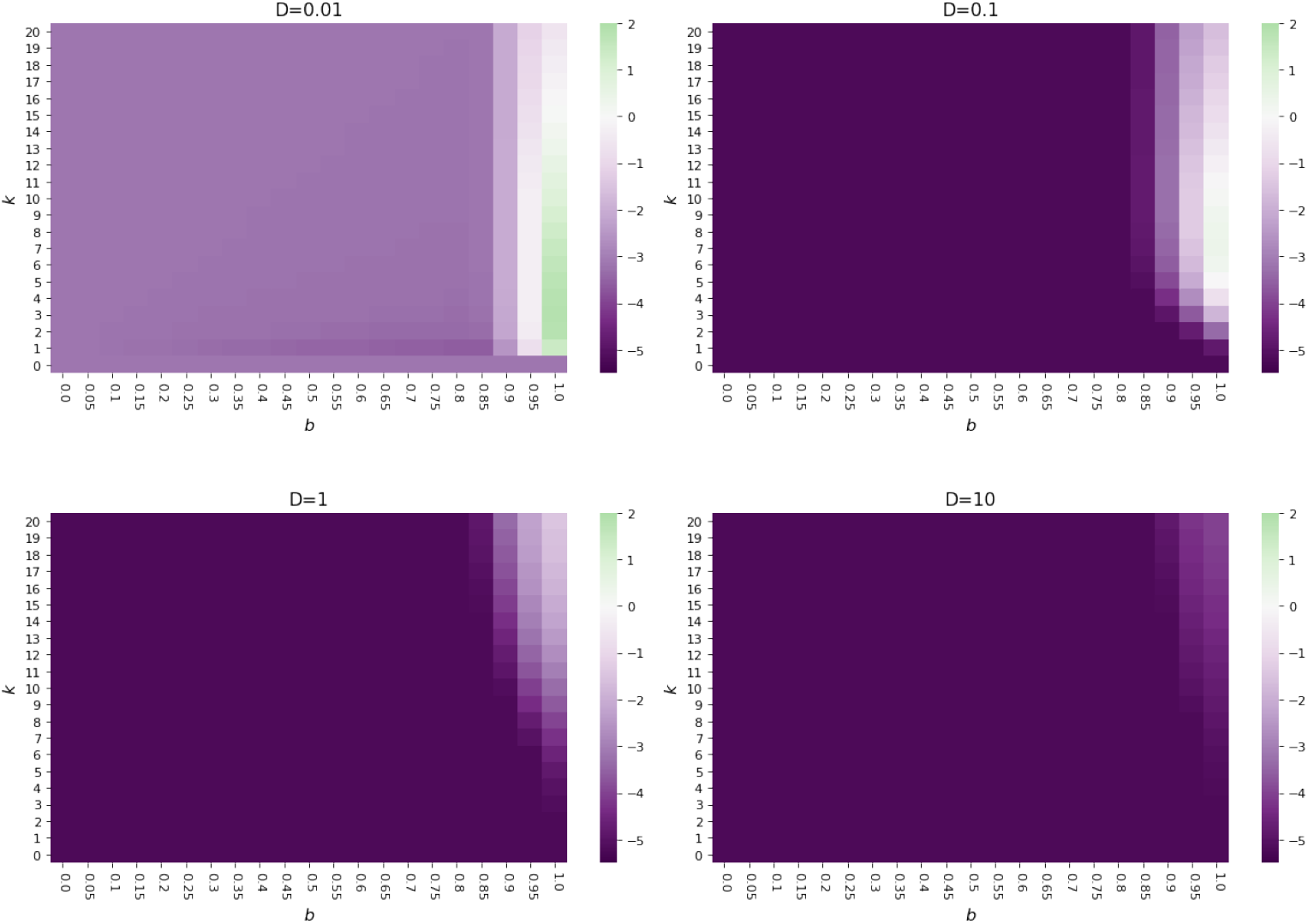
Lower synthesis rate of gene product and no cytoplasmic degradation shifts preferred compartment to nuclear encoding in fluctuating environments. As the bottom row of Figure 3, absolute cost difference between encoding compartments for environmental demand with oscillation amplitude *b* and frequency *k*, for *p* = 0.75, *λ* = 0.1 and *ν*_*c*_ = 0.

**Figure S11:**
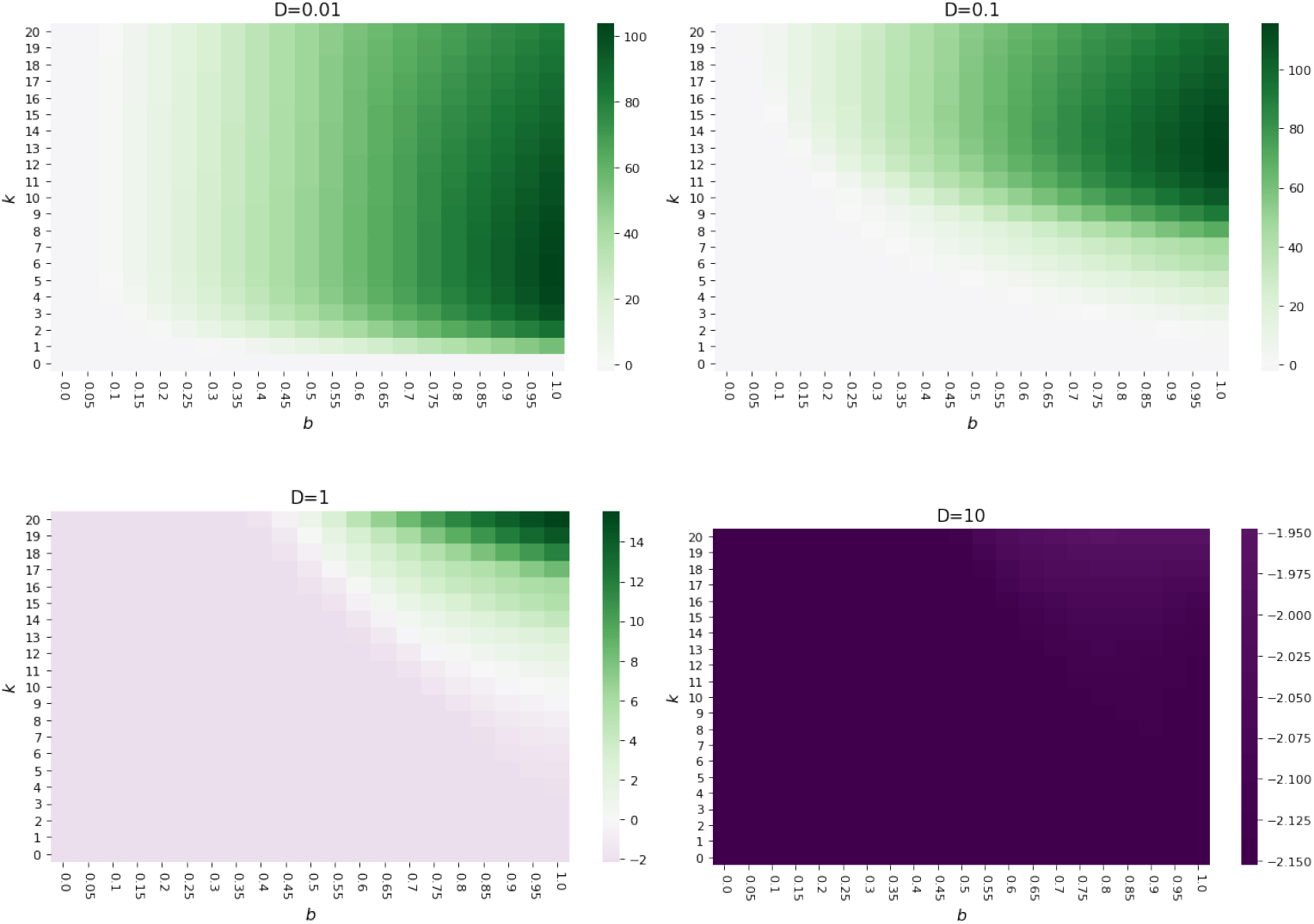
Higher synthesis rate of gene product and no cytoplasmic degradation shifts preferred compartment to organelle encoding in fluctuating environments. As the bottom row of Figure 3, absolute cost difference between encoding compartments for environmental demand with oscillation amplitude *b* and frequency *k*, for *p* = 0.75, *λ* = 10 and *ν*_*c*_ = 0.

**Figure S12:**
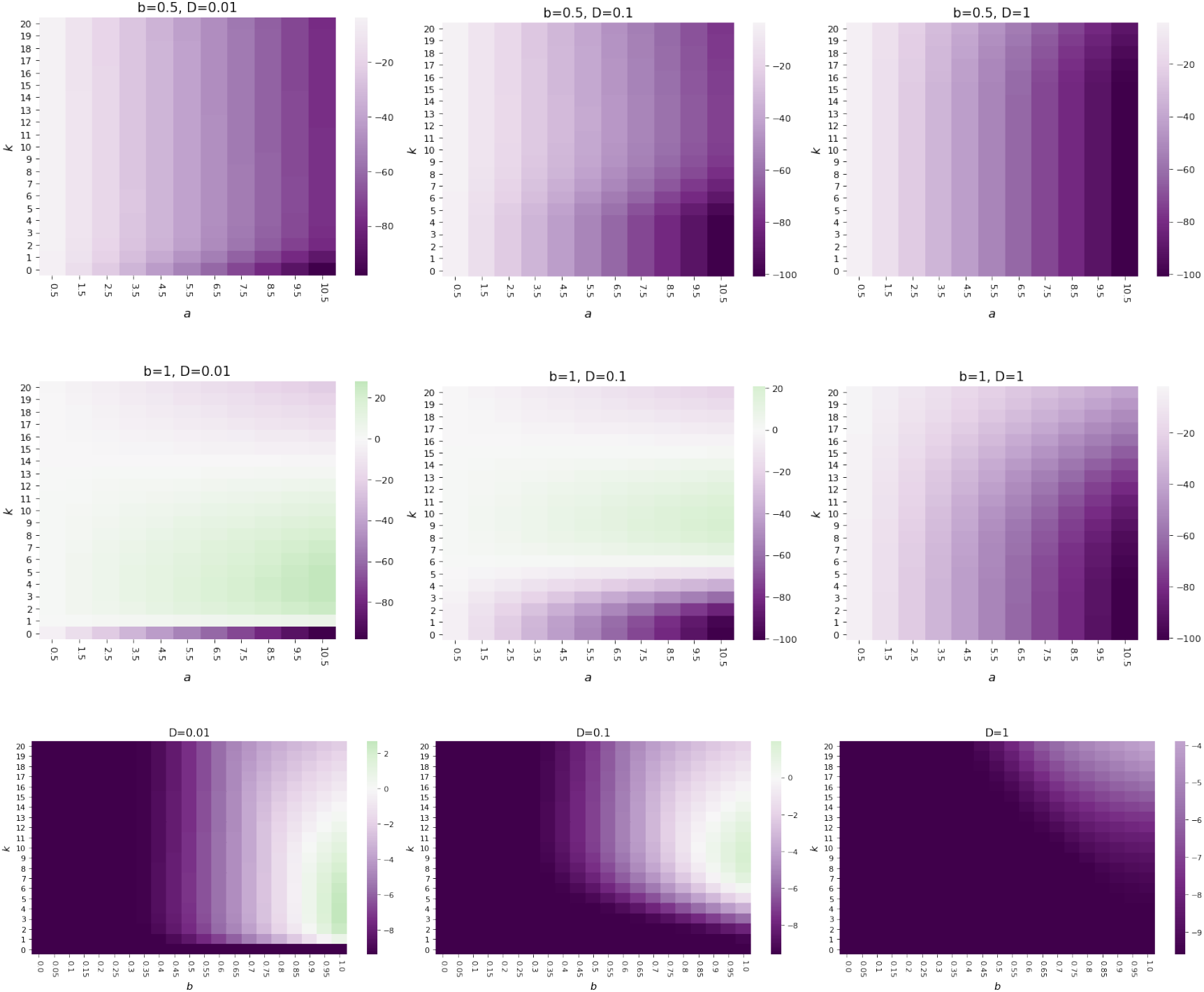
Cost difference for a signalling cost function with the interplay between environmental oscillation and cell biological processes. More dynamical environments favour organelle-encoding and less dynamical environments favour nuclear-encoding. As Figure 3 but defining a cost as the integral over a time interval of the signalling function Eq (1), i.e. 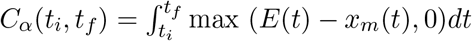.

**Figure S13:**
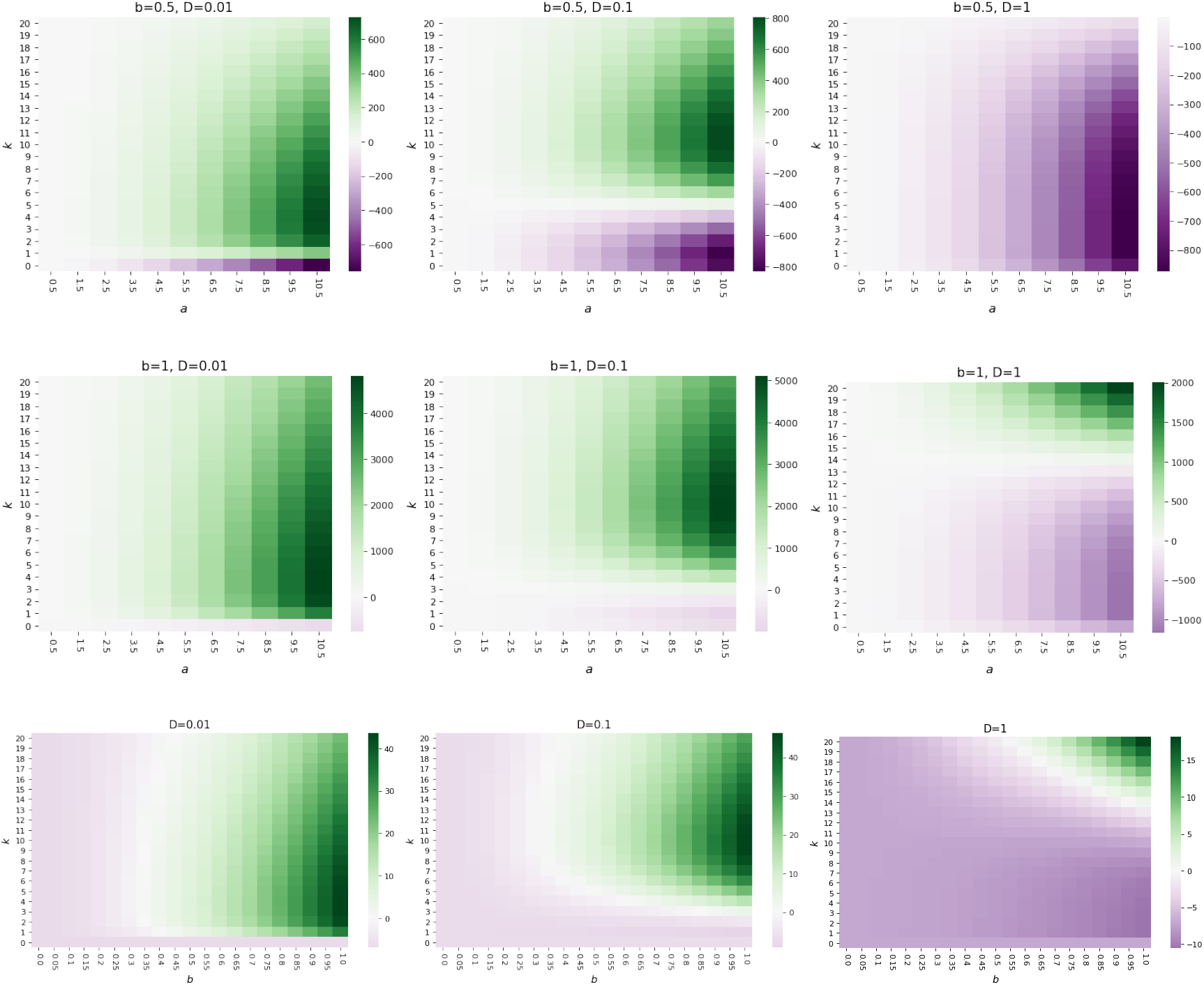
Cost difference for a quadratic cost function with the interplay between environmental oscillation and cell biological processes. More dynamical environments favour organelle-encoding and less dynamical environments favour nuclear-encoding. As Figure 3 but defining a quadratic cost function 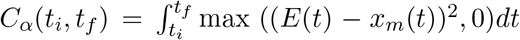. In this case, the signalling function Eq(1) is changed to *f* (*E*(*t*), *x*_*m*_(*t*)) = max ((*E*(*t*) −*x*_*m*_(*t*))^2^, 0). We observe that the organelle-encoding strategy is slightly more favoured for a wider range of amplitudes, including more lower values, for this choice of cost function.

## Notes

https://github.com/StochasticBiology/Environmental-oDNA-retention

### Competing Interest Statement

The authors have declared no competing interest.

